# Clonal hematopoiesis is driven by aberrant activation of TCL1A

**DOI:** 10.1101/2021.12.10.471810

**Authors:** Joshua S. Weinstock, Jayakrishnan Gopakumar, Bala Bharathi Burugula, Md Mesbah Uddin, Nikolaus Jahn, Julia A. Belk, Bence Daniel, Nghi Ly, Taralyn M. Mack, Cecelia A. Laurie, Jai G. Broome, Kent D. Taylor, Xiuqing Guo, Moritz F. Sinner, Aenne S. von Falkenhausen, Stefan Kääb, Alan R. Shuldiner, Jeffrey R. O’Connell, Joshua P. Lewis, Eric Boerwinkle, Kathleen C. Barnes, Nathalie Chami, Eimear E. Kenny, Ruth J. Loos, Myriam Fornage, Lifang Hou, Donald M. Lloyd-Jones, Susan Redline, Brian E. Cade, Bruce M. Psaty, Joshua C. Bis, Jennifer A. Brody, Edwin K. Silverman, Jeong H. Yun, Dandi Qiao, Nicholette D. Palmer, Barry I. Freedman, Donald W. Bowden, Michael H. Cho, Dawn L. DeMeo, Ramachandran S. Vasan, Lisa R. Yanek, Lewis C. Becker, Sharon Kardia, Patricia A. Peyser, Jiang He, Michiel Rienstra, Pim Van der Harst, Robert Kaplan, Susan R. Heckbert, Nicholas L. Smith, Kerri L. Wiggins, Donna K. Arnett, Marguerite R. Irvin, Hemant Tiwari, Michael J. Cutler, Stacey Knight, J Brent. Muhlestein, Adolfo Correa, Laura M. Raffield, Yan Gao, Mariza de Andrade, Jerome I. Rotter, Stephen S. Rich, Russell P. Tracy, Barbara A. Konkle, Jill M. Johnsen, Marsha M. Wheeler, J. Gustav Smith, Olle Melander, Peter M. Nilsson, Brian S. Custer, Ravindranath Duggirala, Joanne E. Curran, John Blangero, Stephen McGarvey, L. Keoki Williams, Shujie Xiao, Mao Yang, C. Charles. Gu, Yii-Der Ida. Chen, Wen-Jane Lee, Gregory M. Marcus, John P. Kane, Clive R. Pullinger, M. Benjamin Shoemaker, Dawood Darbar, Dan Roden, Christine Albert, Charles Kooperberg, Ying Zhou, JoAnn E. Manson, Pinkal Desai, Andrew Johnson, Rasika Mathias, Thomas W. Blackwell, Goncalo R. Abecasis, Albert V. Smith, Hyun M. Kang, Ansuman T. Satpathy, Pradeep Natarajan, Jacob Kitzman, Eric Whitsel, Alexander P. Reiner, Alexander G. Bick, Sidd Jaiswal, on behalf of the NHLBI Trans-Omics for Precision Medicine (TOPMed) Consortium

## Abstract

A diverse set of driver genes, such as regulators of DNA methylation, RNA splicing, and chromatin remodeling, have been associated with pre-malignant clonal expansion of hematopoietic stem cells (HSCs). The factors mediating expansion of these mutant clones remain largely unknown, partially due to a paucity of large cohorts with longitudinal blood sampling. To circumvent this limitation, we developed and validated a method to infer clonal expansion rate from single timepoint data called PACER (passenger-approximated clonal expansion rate). Applying PACER to 5,071 persons with clonal hematopoiesis accurately recapitulated the known fitness effects due to different driver mutations. A genome-wide association study of PACER revealed that a common inherited polymorphism in the *TCL1A* promoter was associated with slower clonal expansion. Those carrying two copies of this protective allele had up to 80% reduced odds of having driver mutations in *TET2, ASXL1, SF3B1, SRSF2*, and *JAK2*, but not *DNMT3A. TCL1A* was not expressed in normal or *DNMT3A*-mutated HSCs, but the introduction of mutations in *TET2* or *ASXL1* by CRISPR editing led to aberrant expression of *TCL1A* and expansion of HSCs in vitro. These effects were abrogated in HSCs from donors carrying the protective *TCL1A* allele. Our results indicate that the fitness advantage of multiple common driver genes in clonal hematopoiesis is mediated through *TCL1A* activation. PACER is an approach that can be widely applied to uncover genetic and environmental determinants of pre-malignant clonal expansion in blood and other tissues.

## Main

Aging is characterized by the accumulation of somatic mutations, nearly all of which are “passengers” that have little fitness consequence on the cells in which they occur. However, infrequent fitness-increasing mutations, called “drivers”, may result in an expanded lineage of cells, termed a clone. Clonal hematopoiesis of indeterminate potential (CHIP) is defined by the acquisition of specific, cancer-associated driver mutations in hematopoietic stem cells (HSC) from persons without a blood cancer^1^. Previous reports have associated CHIP with increased risk for hematologic malignancy, coronary heart disease, and mortality^2–6^. The variant allele fraction (VAF), defined as the proportion of sequencing reads at a locus containing the mutant allele, is an approximate measure of clone size. In contrast to low VAF clones, which are ubiquitous in older individuals^7^, large VAF CHIP clones are less common and more likely to result in hematologic malignancy and cardiovascular disease^5,6,8,9^.

The genes commonly mutated in CHIP include regulators of DNA methylation (*TET2, DNMT3A*), chromatin remodeling (*ASXL1*), and RNA splicing (*SF3B1, SRSF2, U2AF1*). Even though these mutations are highly prevalent in CHIP and hematological cancers, the mechanisms driving clonal expansion remain largely unknown. This is partially due to a lack of sizable cohorts with serially sampled blood over decades which would otherwise enable studies on genetic and environmental correlates of clonal expansion. To address this gap, we developed a method for approximating the rate of clonal expansion from a single timepoint, termed PACER, which was validated using longitudinal sequencing over 10 years in 55 CHIP carriers. We then used PACER to perform the first large-scale investigation of the germline determinants of clonal expansion in 5,071 CHIP carriers from the NHLBI Trans-Omics for Precision Medicine (TOPMed) program^10,11^, which revealed activation of *TCL1A* as an event driving clonal expansion for multiple mutated genes in CHIP.

### Derivation and validation of PACER

We identified high-confidence somatic mutations in peripheral blood DNA by analyzing TOPMed whole genome sequencing (WGS) data with Mutect2^12^. To remove sequencing artifacts and germline variants we performed stringent variant filtering and quality control. We identified CHIP mutations in 5,071 individuals using a curated list of leukemogenic driver mutations (Supplemental Table 1) (Methods). As described in our previous report^11^, the prevalence of CHIP was strongly associated with age at blood draw, and >75% of these mutations were in *DNMT3A, TET2*, or *ASXL1*.

In HSCs, passenger mutations accrue at a rate that is fairly constant over time and that is similar across individuals^13–15^. Thus, the number of passenger mutations in the founding cell of a CHIP clone can be used to approximate the date of acquisition of the driver mutation (Figure 1a). Prior studies have enumerated passenger mutation burden in HSCs by performing WGS on colonies derived from single cells^16,17^. We theorized that the passenger mutation burden in the founding cell for a CHIP clone could instead be approximated from WGS of whole blood DNA without isolation of single cells. As a mutant clone expands, the VAF of both the driver and passenger mutations increases. The number of passengers in any given cell is simply the sum of the mutations present prior to the acquisition of the driver event (ancestral passengers) and mutations acquired after the driver event (sub-clonal passengers). Because the limit of detection for mutations from WGS at ∼38X coverage depth is ∼8-10% VAF, the detectable passengers in whole blood DNA are far more likely to be ancestral passengers than sub-clonal passengers. This is because the sub-clonal passengers are private to each subsequent division of the original mutant cell, and, in the absence of a second driver event, quickly fall below the limit of detection in WGS data from bulk tissue (Supplementary Text 1). Furthermore, as the size of the clone also determines the number of detectable passengers from WGS due to the limited sensitivity of detection at 38X depth, high fitness clones will harbor more detectable passengers than lower fitness clones that arose at the same time. Based on these observations, we used the detectable passengers as a composite measure of clone fitness and birth date. For two individuals of the same age and with clones of the same size, we expect the clone with more passengers to be more fit, as it must have expanded to the same size in less time.

**Fig 1.**
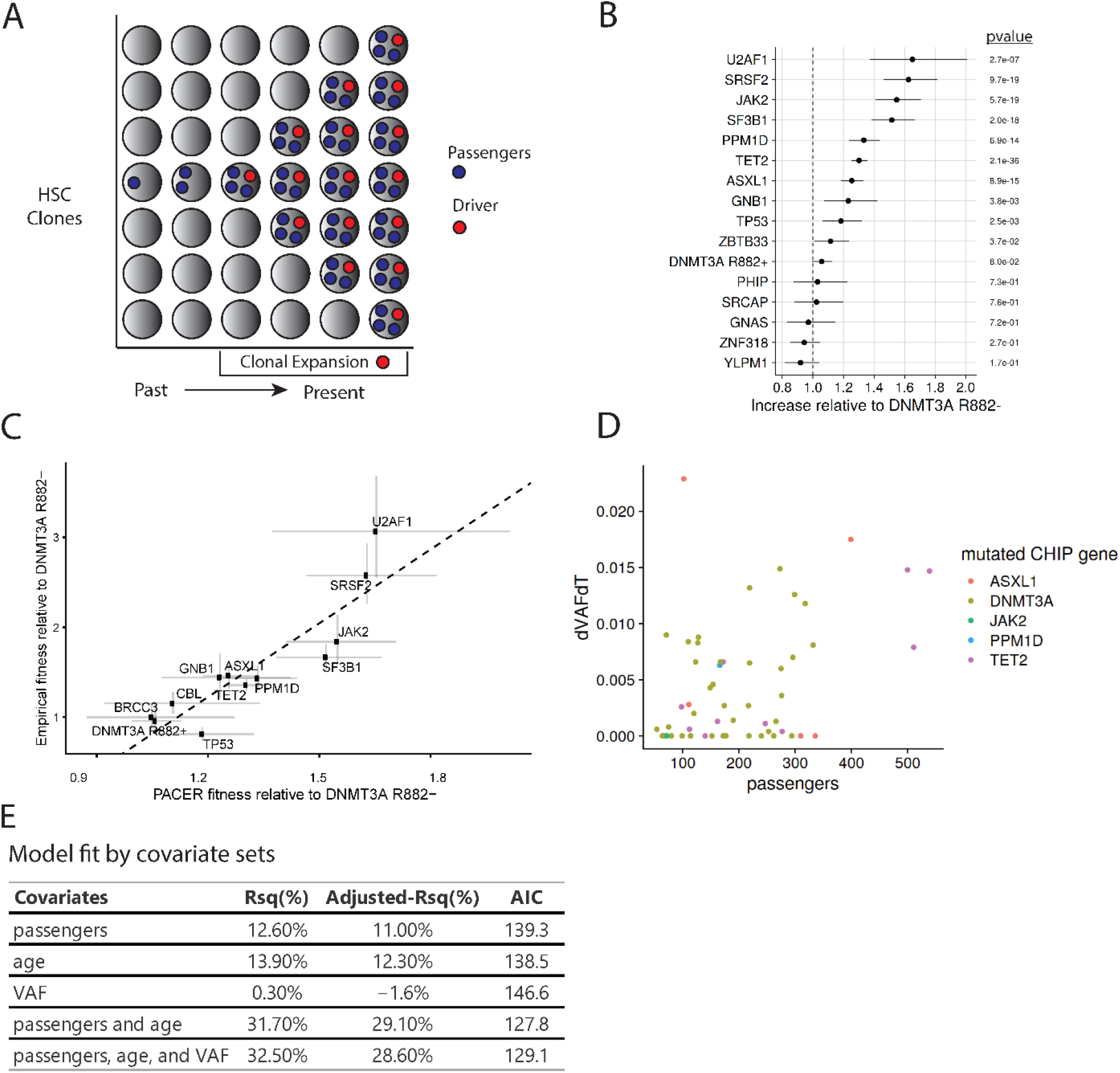
PACER Enables Estimation of Clonal Expansion from a Single Blood Draw. **A**, A schematic depiction of using passenger counts to estimate the rate of expansion of a hematopoietic stem cell (HSC) clone after the acquisition of a driver mutation. The passengers (blue) that precede the driver (red) can be used to date the acquisition of the driver. **B**, The relative abundances of passenger counts were estimated for CHIP driver genes with at least 30 cases using a negative binomial regression, adjusting for age at blood draw, driver VAF, and study. The coefficients are relative to DNMT3A R882-CHIP. **C**, The relative abundances of passenger counts are plotted against the empirical estimates of gene fitness derived from the longitudinal deep sequencing in Fabre et al.^17^. The estimate of the association from weighted least squares (slope = 2.7, pvalue = 9.6 × 10^−5^, R^2^ = 80%) is plotted as a dashed line. **D**, The observed clonal expansion rates (dVAFdT), as expressed in the change in variant allele frequency (VAF) over time (years), were associated with increased passenger counts in 55 CHIP carriers from the Women’s Health Initiative. Colors indicate the mutated driver gene. **E**, A multivariate model including passenger counts, age at blood draw, and VAF indicates the relative contributions of age and VAF over baseline models. AIC is Akaike information criteria, where smaller values indicate better model fit.

To estimate the number of passenger mutations, we first performed genome-wide somatic variant calling for 5,071 CHIP carriers and 23,320 controls without CHIP driver mutations. As these raw variant calls contain a combination of true somatic variants, germline variants, and sequencing artifacts, we implemented a series of stringent filters to enrich for the detection of true passengers (see Methods). We first selected only those variants that were found in a single individual in the dataset, as recurrent variants are enriched for germline polymorphisms and recurrent artifacts. We also excluded variants with a VAF greater than 35%, as these would also be enriched for germline polymorphisms. Since different base substitutions varied in their association with age at blood draw, we selected only C>T and T>C mutations, as these were the most strongly age-associated in our data, consistent with prior work identifying such mutations as essential elements of the “clock-like” signature^18^.

Amongst the 5,071 CHIP carriers, individuals had on average 271 passengers in WGS identified by our approach (interquartile range: 142 – 317). The passengers were increased by 54% (95% CI: 51%-57%) in the CHIP carriers (Extended Data Fig 1) compared to the controls after adjusting for age and study using a negative binomial regression. In the controls without CHIP, we presumed the detected passengers were reflective of clonal hematopoiesis without known driver mutations or due to drivers we could not assess such as mosaic chromosomal alterations (mCAs)^19^. Some of these could also have been incompletely removed artifacts. The passengers were also positively associated with age, on average increasing by 13.7% (95% CI: 13.0%-14.3%) each decade. Of the CHIP carriers in TOPMed, 89% had a single driver mutation. We found that each additional driver mutation detected in a given sample was associated with an increase in passenger mutation counts (Extended Data Fig 2). This is likely due to the presence of cooperating driver mutations in the same clone in these persons, as each successive expansion caused by a new driver mutation captures additional passenger mutations that accumulated in the time between the last driver event and the newer one. For this reason, we limited further analyses on clonal expansion rate only to the 4,536 CHIP carriers with a single driver event.

We validated the passengers as an estimator of fitness both theoretically and empirically. For the theoretical validation, we constructed a simulation of HSC dynamics to characterize the relationship between fitness and detectable passenger counts (Supplementary Text 1). The simulation indicated that founding passengers were associated with driver fitness (spearman *ρ* =0.09, pvalue < 2 × 10^−16^). We estimated a passenger mutation rate per diploid genome per year of 2.3, or a per-base pair rate of 3.83 × 10^−10^. Assuming 100,000 HSCs^14,20^, this results in a per-base-pair passenger mutation rate of 3.83 × 10^−15^ per HSC clone per year without correction for the sensitivity of the sequencing technology used. This number is substantially lower than previous estimates using WGS from single hematopoietic colonies, likely due the low sensitivity of detecting true passengers in whole blood DNA compared to the gold standard of single-cell derived colonies and also because we limited the base substitutions in our analysis to C>T or T>C. Nonetheless, we were able to use these data to derive a hierarchical Bayesian estimator of clone fitness (Methods), which adjusts for age at blood draw and cohort effects, and confirmed its correspondence to the observed passenger counts.

### PACER predicts fitness of distinct driver mutations

Building on recent computational estimates of variant fitness^20^, we estimated the distribution of passenger counts for the most common CHIP driver genes. We used non-R882 *DNMT3A* mutations as a reference point and estimated the relative abundances of passengers in other genes using negative binomial regression adjusting for age, VAF, and study. We termed this approach PACER (passenger-approximated clonal expansion rate). Mutations in splicing factors (*SF3B1, SRSF2, U2AF1*) and *JAK2* V617F mutations were the fastest growing according to PACER, while *DNMT3A* R882-was among the slowest (Figure 1b-c, Supplementary Table 2). Mutations in *TET2, ASXL1, PPM1D, TP53, ZBTB33*, and *GNB1* were in the next tier and had approximately the same level of fitness estimated from *PACER*. Relative to the R882-carriers, we observed a modest increase in fitness in *DNMT3A* R882 mutant clones. These observations are concordant with prior empirical estimates of variant fitness derived from longitudinal sequencing of samples with clonal hematopoiesis^6,17,21,22^ and provides further validation of our approach.

To empirically validate the predictive ability of passenger count, we performed targeted sequencing for driver variants from two blood samples taken approximately 10 years apart in 55 CHIP carriers from the Women’s Health Initiative (WHI, Methods). WGS from the first time point was used to determine passenger count. We quantified clonal expansion by dividing the change in VAF by the change in time (years) 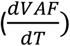 of the driver variants identified at the first blood draw. Of the sequenced carriers, 40 had clones with a single CHIP mutation that were constant in size or expanded. We constructed a simple estimator of 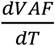 using only the passengers, VAF, and age from the first blood draw (Methods). Our theoretical framework considered passengers to be an estimate of clone fitness after accounting for age and VAF, hence these latter two variables were also considered in the model. A model only including VAF had lower predictive ability (Rsq = 0.30%, Adjusted Rsq = -1.60%) for clonal expansion than a model only including passengers (Rsq = 12.6 %, Adjusted Rsq = 11%). A model including only age had similar performance (Rsq = 13.9%, Adjusted Rsq = 12.3%) to the passenger model. A model that included age and VAF in addition to passenger count improved the prediction of clonal expansion (Rsq = 32.5%, Adjusted Rsq = 28.6%, Figure 1d-e). These results suggested that inferring clonal expansion from age- and VAF-adjusted passenger mutation counts was able to not only describe past growth, but also predict future growth rate.

### Genome wide association study identifies inherited determinants of clonal expansion

We performed a genome-wide association study (GWAS) of PACER in CHIP carriers to identify inherited genetic variation that associates with clonal expansion. Association analyses were performed using the SAIGE^23^ statistical package. We included age at blood draw, study, VAF, and the first ten genetic ancestry principal components as covariates.

The GWAS identified a single locus at genome-wide significance overlapping *TCL1A* (Figure 2a). We used SuSIE^24^ to perform genetic fine-mapping to identify the most likely causal set of variants, which further narrowed down the associated region to a credible set containing a single variant, rs2887399 (Extended Data Fig. 3). Each additional alternative (alt) allele (T) was associated with a 0.15 decrease in passenger count z-score (pvalue = 4.5 ×10^−12^). The alt-allele is common, occurring in 26% of haplotypes sequenced in TOPMed. rs2887399 lies in the core promoter of *TCL1A* as defined by the Ensembl regulatory build^25^, 162 base-pairs from the canonical transcription start site (TSS) and in a CpG island. Analysis of the variant by the Open Targets^26^ variant-to-gene prediction algorithm also nominated *TCL1A* as the causal gene. We did not find any association between PACER and rare variants near rs2887399, suggesting that rs2887399 is not tagging other genetic variants and is the causal variant at this locus (Extended Data Fig. 4-5). *TCL1A* has been implicated in lymphoid malignancies as a translocation partner in T-prolymphocytic leukemia^27^, but it has not been studied in the context of HSC biology. *TCL1A* is also the only gene in the duplicated region of chromosome 14q32 associated with an inherited predisposition to develop myeloid malignancies shared by all kindreds^28,29^. Of note, the region in the *TCL1A* promoter where rs2887399 resides is only partially conserved between humans and other primates, and poorly conserved with non-primate species (Extended Data Fig 6).

**Fig 2.**
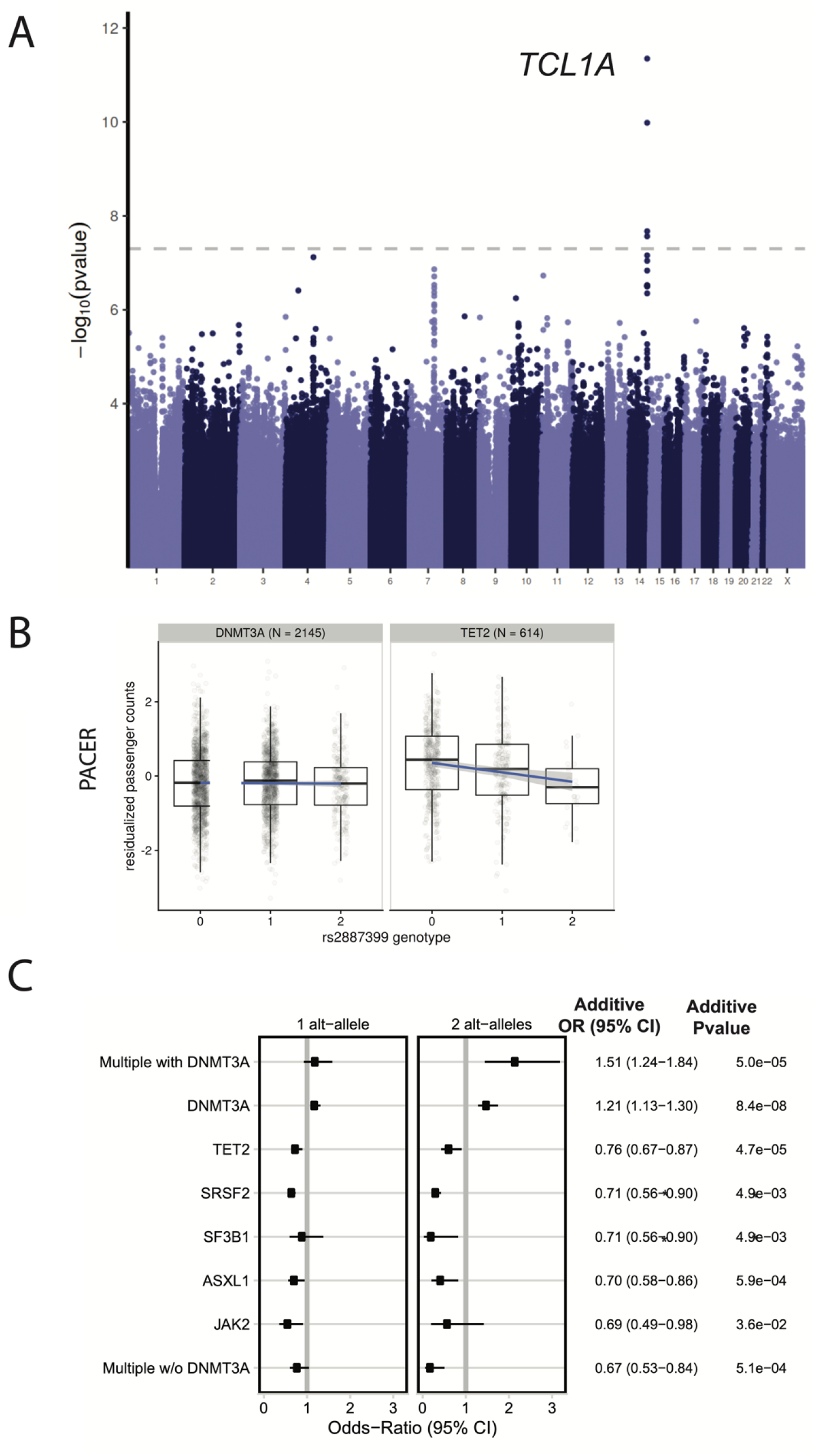
GWAS of PACER Identifies Germline Determinants of Clonal Expansion in Blood. **A**, A genome-wide association study (GWAS) of passenger counts identifies *TCL1A* as a genome-wide significant locus. **B**, The association between the genotypes of rs2887399 and PACER varied between *TET2* and *DNMT3A*. Alt-alleles were associated with decreased PACER score in *TET2* mutation carriers, in contrast to *DNMT3A* carriers, where no association was observed. **C**, The association between alt-alleles at rs2887399 and presence of specific CHIP mutations varies by CHIP mutations. Forest plot shows the effect estimates of a single T allele and two T-alleles respectively, estimating using Firth logistic regression. On the right of the forest plot, effect estimates and p-values are included from SAIGE^23^, which uses an additive coding of the alt-alleles for hypothesis testing. In the additive tests, *SF3B1* and *SRSF2* were grouped together to aid convergence.

We next performed a genome-wide search of rare variation associated with the passengers. We identified 15 windows associated with passenger counts at Bonferroni significance (pvalue = 2.9 × 10^−5^, Supplementary Table 4). We identified an intergenic region 113kb from the TSS of *TNFAIP3* (pvalue = 5.4 × 10^−7^) that is a distal enhancer of *TNFAIP3* (GeneHancer^30^).

### Association of *rs2887399* with specific driver genes

We asked whether the association between rs2887399 and PACER was modified by CHIP driver gene. Using *DNMT3A* as the reference, we investigated whether other genes had different effect estimates for rs2887399. We observed that alt-allele dosage in rs2887399 was more protective against clonal expansion in *TET2* than *DNMT3A* (beta = -0.24, pvalue = 9.6 × 10^−4^, Figure 2b, Supplementary Table 5).

Clones with a decreased expansion rate may never grow large enough to be detected, so we also performed association tests between rs2887399 and presence of a CHIP-associated driver mutation stratified by gene. In our previous analysis^11^, we reported that the alt-allele was associated with increased risk for *DNMT3A* mutations. Prior reports have also identified that the alt-allele of rs2887399 decreases risk for mosaic loss of the Y chromosome (LOY)^31^ (OR = 0.80, pvalue = 4.3 ×10^−136^). Here, we observed that rs2887399 was associated with significantly reduced odds of mutations in *TET2, ASXL1, SF3B1, SRSF2*, and possibly *JAK2* (Figure 2c). The effect size of rs2887399 was large for a common variant, as those carrying 2 copies of the alt-allele had odds ratios for having a driver mutation in these genes ranging from 0.22 to 0.63 (Figure 2d). The risk reduction was particularly strong for mutations in *SF3B1* and *SRSF2*, as well as for having >1 non-*DNMT3A* driver mutations (Figure 2c-d, Supplementary Table 6-7, Methods). The latter group is particularly relevant clinically, as these persons have a high risk of risk of transformation, and in some cases may already have early-stage MDS^5,6,32^. In sum, these results indicate that the alt-allele at rs2887399 is protective against CHIP due to driver mutations in several genes that have higher risk of progression to frank hematologic malignancy^6^.

Previous analyses in UK Biobank^33^ have also implicated rs2887399 in reduced blood cell counts (Supplementary Table S8), consistent with an effect on hematopoiesis, but it is unknown if this is independent of hematological malignancy or CHIP.

### *TCL1A* expression in hematopoietic cells

Next, we sought to establish how rs2887399 might shape the hematologic phenotypes observed. We first asked if the variant was associated with *TCL1A* expression in any cell type. As identified in the GTEx v8 eQTL release^34^, the alt-allele reduces expression of *TCL1A* in whole blood (normalized effect size = -0.13, pvalue = 1.4 × 10^−5^). The GWAS of PACER colocalized with cis-expression quantitative trait loci (eQTLs) for *TCL1A* in whole blood (posterior probability of a single shared causal variant = 97.1%, Extended Data Fig 7). The association in whole blood is likely driven by B-cells, as *TCL1A* is highly expressed in B-cells but appears to have absent or low expression in all other cell types in blood except for rare plasmacytoid dendritic cells (Supplementary Table 9, Supplementary Figure 1).

Little is known about *TCL1A* expression in HSCs. We examined whether CHIP-associated mutations altered the regulation of the *TCL1A* locus in human hematopoietic stem and progenitor cells (HSPCs) using publicly available single-cell RNA sequencing (scRNAseq) and ATAC-sequencing (ATAC-seq) datasets of normal and malignant hematopoiesis. *TCL1A* was expressed in fewer than 1 in 1000 cells identified as HSC/MPPs in scRNAseq data from 6 normal human marrow samples (range 0-0.17%)^35,36^. In contrast, *TCL1A* was expressed in a much higher fraction of HSC/MPPs in 3 out of 5 samples from persons with *TET2* or *ASXL1*-mutated myeloid malignancies (range 2.7-7%) (Figure 3a). Next, using a dataset of ATAC-seq in normal and pre-leukemic HSCs (pHSCs)^29^, we evaluated chromatin accessibility at the *TCL1A* promotor. Consistent with the lack of *TCL1A* transcripts in normal HSCs, we observed that the promoter was not accessible in either normal human donor HSCs or in HSCs from patients with AML that were not part of the mutant clone. We also did not observe accessible chromatin in two carriers of *DNMT3A* mutated pHSCs. In contrast, the two patients with *TET2* mutated pHSCs had clearly accessible chromatin at the *TCL1A* promoter (Figure 3b).

**Figure 3.**
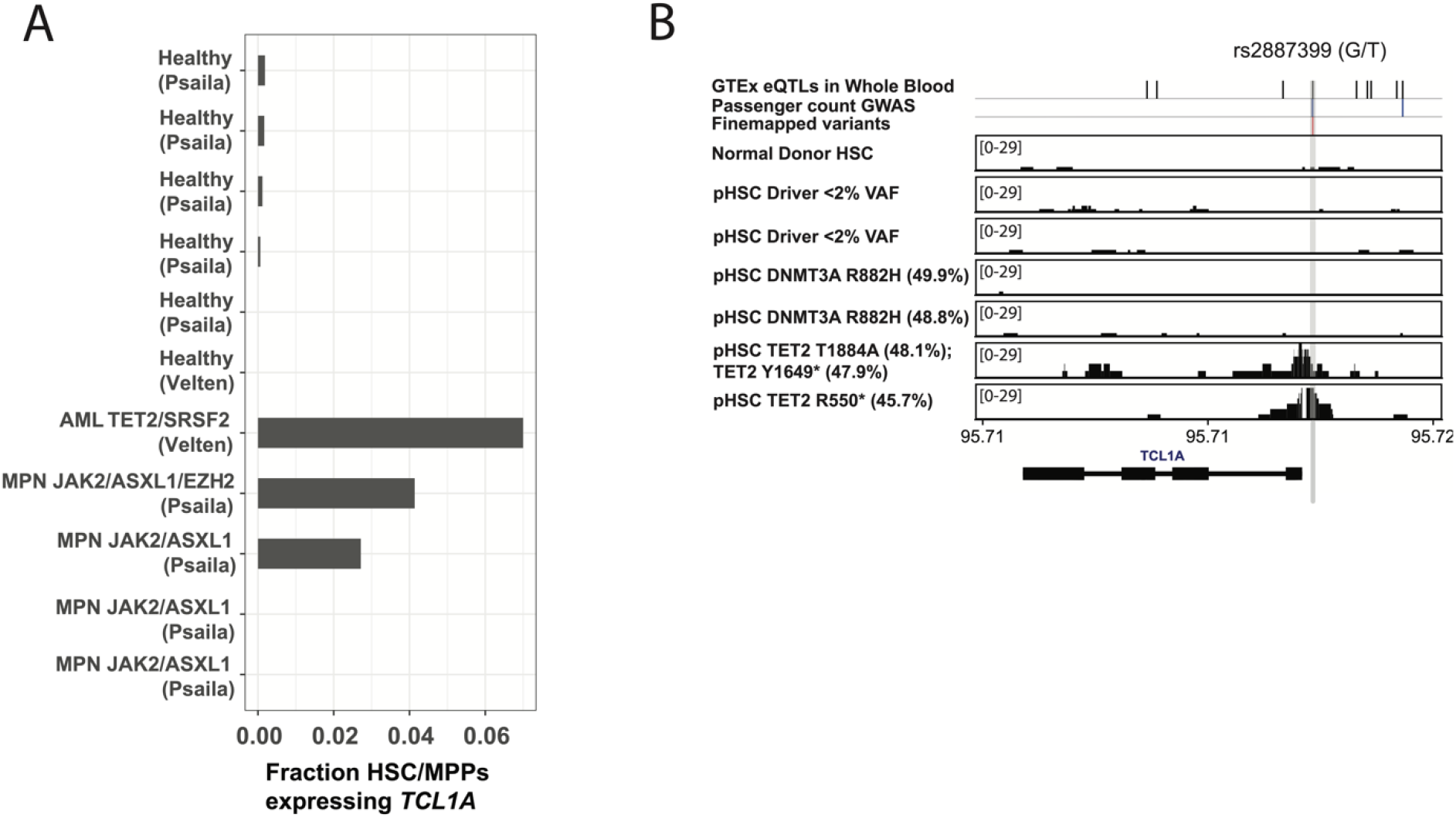
*TET2* and *ASXL1* mutations permit aberrant *TCL1A* accessibility and transcript expression in HSCs and MPPs. **A**. Quantification of fraction of HSCs and MPPs expressing *TCL1A* transcripts in patients with *TET2* or *ASXL1* driven acute myeloid leukemia (AML) or myeloproliferative neoplasm (MPN) compared to healthy donors. Data is from single-cell RNA sequencing generated in Psaila et al^36^and Velten et al^35^. **B**. ATAC-sequencing tracks of the *TCL1A* locus near rs2887399 in HSCs form healthy donors (row 1), pre-leukemic hematopoietic stem cells (pHSCs) from patients with AML but no detected driver mutations (rows 2-3), pHSCs with *DNMT3A* mutations (rows 4-5), and in pHSCs with *TET2* mutations (rows 6-7). Amino acid change and variant allele fraction (VAF) for the driver mutations are shown. Data is from Corces et al^65^. Vertical grey bar indicates location of the rs2887399 SNP. Black hash marks indicate positions of GTEX v8 eQTLs for TCL1A in whole blood, blue hash marks indicate positions of genome-wide significant SNPs, and the red hash mark indicates the position of the single causal variant identified by fine-mapping, rs2887399.

### Functional effect of rs2887399 on normal and CHIP-mutated HSCs

These observations led us to propose the following mechanistic model: Normally, the *TCL1A* promoter is inaccessible and gene expression is absent or very low in HSCs. In the presence of driver mutations in *TET2, ASXL1, SF3B1, SRSF2*, or LOY, *TCL1A* is aberrantly expressed and drives clonal expansion of the mutated HSCs. The presence of the alt-allele of rs2887399 inhibits accessibility of chromatin at the *TCL1A* promoter, leading to reduced expression of *TCL1A* RNA and protein and abrogation of the clonal advantage due to the mutations (Extended Data Fig 8).

To test our model experimentally, we first obtained human CD34+ mobilized peripheral blood cells from donors who were GG (homozygous reference), TT (homozygous alternate), or GT (heterozygous) genotype at rs2887399. The three donors were healthy and between 29-32 years old at the time of donation. To mimic CHIP-associated mutations, we used CRISPR to introduce insertion-deletion mutations in *DNMT3A, TET2*, or *ASXL1* in HSPCs for each rs2887399 genotype. Editing at the adeno-associated virus integration site 1 (AAVS1) was done as a control for each rs2887399 genotype (Figure 4a). High efficiency of editing was confirmed by Sanger sequencing (Extended Data Fig 9).

**Figure 4.**
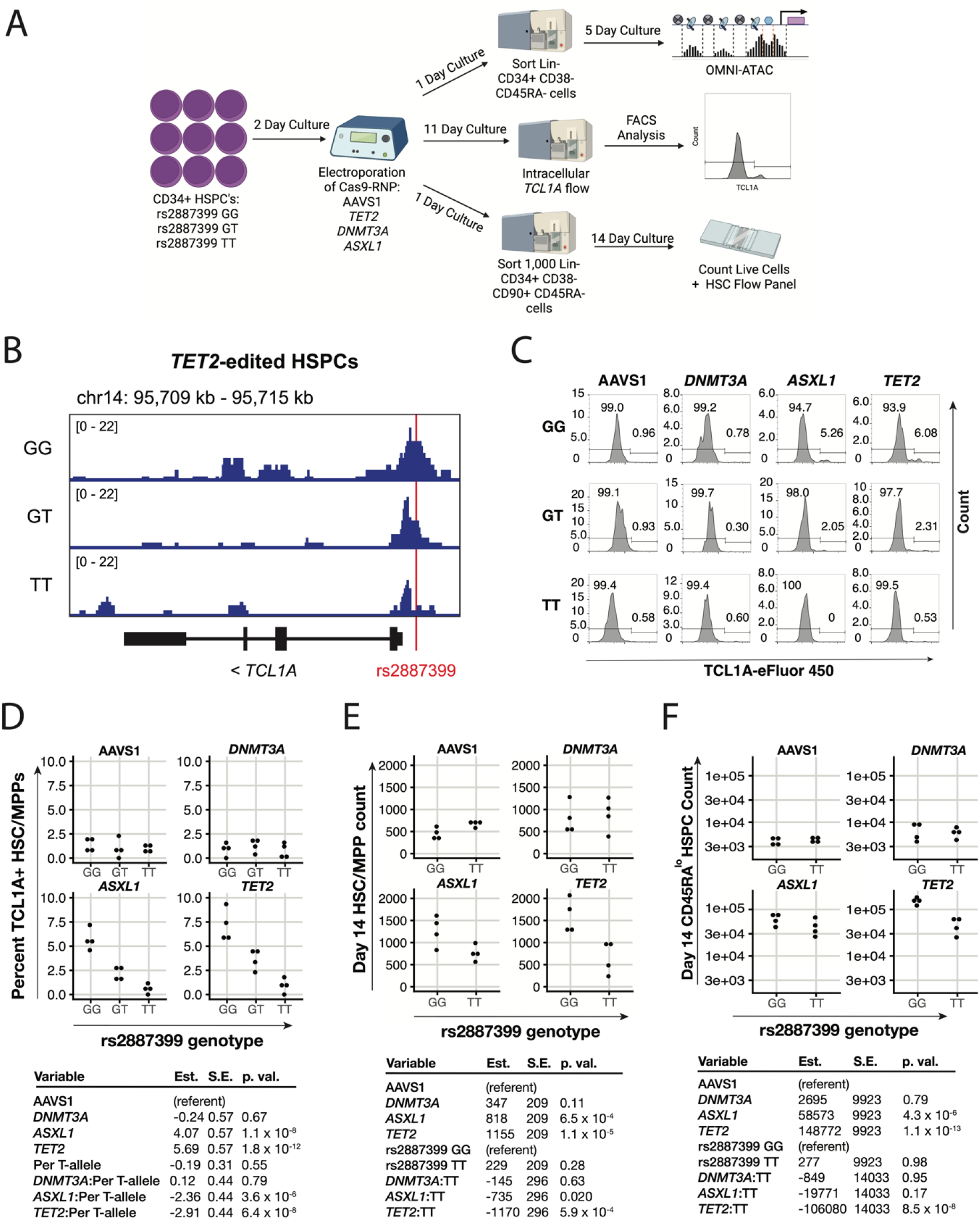
T allele of rs2887399 reduces TCL1A expression and extinguishes clonal expansion phenotype of *TET2* and *ASXL1* mutant HSPCs. **A**. Schematic of experimental workflow. Human HSPCs from donors carrying rs2887399 GG, GT, or TT genotypes were electroporated with Cas9 targeting AAVS1, *TET2, DNMT3A*, or *ASXL1* and cultured for OMNI-ATAC, intracellular flow cytometric analysis of TCL1A expression, or an *in vitro* HSPC expansion assay. **B**. ATAC-sequencing tracks illustrating chromatin accessibility at rs2887399 in *TET2*-edited HSPCs cultured for 5 days from donors of the GG, GT, and TT genotypes. Red line indicates location of rs2887399. **C**. Representative intracellular flow plots of TCL1A protein expression in edited HSCs/MPPs from each rs2887399 donor after 11 days in culture. **D**. Quantification of percent HSCs/MPPs expressing TCL1A from flow cytometry, stratified by edited gene and rs2887399 genotype. Results of a linear regression model for the effect of edited gene (referent to AAVS1), number of T-alleles at rs2887399, and the interaction term of edited gene with T-alleles are presented below. Est. = estimate, S.E. = standard error, p. val. = p-value. **E**. Quantification of Lin-CD34+ CD38-CD45RA-HSC/MPP counts after 14 days of *in vitro* expansion stratified by edited gene and rs2887399 genotype. Results of a linear regression model for the effect of edited gene (referent to AAVS1), rs2887399 genotype (referent to GG), and the interaction term of edited gene with rs2887399 genotype are presented below. **F**. Quantification of Lin-/lo CD34+ CD38-CD45RAlo HSPCs (CD45RAlo HSPCs) after 14 days of *in vitro* expansion stratified by edited gene and rs2887399 genotype. Results of a linear regression model for the effect of edited gene (referent to AAVS1), rs2887399 genotype (referent to GG), and the interaction term of edited gene with rs2887399 genotype are presented below.

First, we examined whether the accessibility of the *TCL1A* promoter seen in the setting of *TET2* mutations was altered by rs2887399 genotype. We edited bulk CD34 cells from each genotype for *TET2*, sorted cells with a marker profile of HSCs and multipotent progenitors (MPPs) (Lineage-CD34+ CD38-CD45RA-), cultured them for 5 days in cytokine-supported media, and then performed ATAC-seq (Extended Data Fig 10). Consistent with the pre-leukemic HSC data, we detected accessibility at the *TCL1A* promoter in *TET2*-edited cells from the rs2887399 GG donor. However, accessibility at the *TCL1A* promoter was decreased in the *TET2*-edited cells in samples from carriers of the T allele in a dose-dependent manner, indicating that the protective effect of the alt-allele of rs2887399 is mediated by blocking promoter accessibility (Figure 4b). These results also suggest that alterations in the chromatin profile of HSPCs can occur within days after the introduction of a *TET2* mutation.

Next, we asked if the differential chromatin accessibility due to rs2887399 altered TCL1A protein expression in HSCs/MPPs. We edited CD34+ cells from donors with the three rs2887399 genotypes at AAVS1, *DNMT3A, TET2*, and *ASXL1*. After 11 days in culture, we performed a flow cytometry-based assay for TCL1A protein expression. We found that ∼1% of HSCs/MPPs from AAVS1 or *DNMT3A* edited samples were positive for TCL1A, which did not vary by rs2887399 genotype. In contrast, 4.6-9.3% of HSCs/MPPs from the GG donor that had been edited for *ASXL1* or *TET2* expressed TCL1A, and the proportion of TCL1A positive HSC/MPPs decreased in donor samples with each additional T allele (4 biological replicates per condition) (Figure 4c-d). There was minimal expression of TCL1A in any non-HSC/MPP CD34+ population in any of the samples. Notably, the proportion of TCL1A expressing HSC/MPPs was less than 10% in all samples even though the proportion of mutant cells was >90% (Extended Data Fig 9). This suggests that even in the presence of driver mutations in *TET2* or *ASXL1*, only a fraction of HSC/MPPs are capable of expressing TCL1A at any given time and is consistent with the single-cell RNA sequencing data from hematological malignancy samples (Figure 3b).

Finally, we asked if rs2887399 had any effect on expansion of HSPCs in vitro. For this experiment, we edited the CD34+ cells from GG and TT donors, sorted HSCs (Lin-CD34+ CD38-CD45RA-CD90+), and allowed the cultures to grow for 14 days, at which time cells were counted and analyzed for HSPC markers by flow cytometry. There was a notable expansion of cells bearing markers of HSCs/MPPs in the *ASXL1* and *TET2* edited samples from the rs2887399 GG donor compared to the AAVS1 edited sample, but this effect was abrogated in edited samples from the rs2887399 TT donor. A population of cells that was Lin-/lo CD34+ CD38-CD45RA dim (CD45RA^dim^ HSPCs), presumably progenitors descended from the HSC/MPP population, was also markedly expanded in the *ASXL1* and *TET2* edited samples from the GG donor, but the degree of expansion was partially reversed in the edited samples from the TT donor. There were no differences in any populations in the AAVS1 or *DNMT3A* edited samples based on rs2887399 genotype (4 biological replicates per condition) (Figure 4e-f). Thus, carrying the alt-allele of rs2887399 abrogates the clonal expansion of HSPCs with *ASXL1* and *TET2* mutations in an experimental system.

## DISCUSSION

Here, we have developed a novel method that allows us to infer clonal expansion rate from a single time point. Our results extend and apply recently developed theory on the evolutionary fitness of clones to permit estimation of fitness within a single individual^20^. Unlike prior methods which used the VAFs of driver variants to estimate fitness^20^, our development of a fitness estimator based on passenger mutations counts permits us to perform association tests for other factors associated with clonal expansion, such as inherited genetic variation and environmental exposures.

We performed the first ever GWAS for determinants of clonal expansion rate and identified a common variant of large effect in the promoter of *TCL1A* as the top hit. Remarkably, this single variant, which has previously been linked to reduced risk of LOY^37^, was also associated with protection from driver mutations in *TET2, ASXL1, SF3B1, SRSF2*, and possibly *JAK2*. We also demonstrated with experimental work that *TCL1A* was normally lowly expressed in HSCs, but that the introduction of mutations in *TET2* or *ASXL1* led to expression of the protein, possibly by permitting promoter chromatin accessibility and hence transcription of the gene. This was completely prevented by the alt-allele of rs2887399, explaining the reduction in predicted clonal expansion rate by PACER and decreased prevalence of these driver mutations in those carrying the allele. To our knowledge, *TCL1A* itself is not somatically mutated in CHIP, perhaps because gain-of-function point mutations are not directly possible. How *TCL1A* expression causes clonal expansion of HSCs is an important question for future studies, but could be related to its reported role in AKT activation^38^. Importantly, our results suggest that pharmacologically targeting TCL1A may suppress growth of CHIP and hematological cancers associated with mutations in these genes.

The large protective effect seen with rs2887399 suggests that *TCL1A* expression is likely a dominant factor mediating clonal expansion due to these mutations. This was especially the case for driver mutations in *SRSF2* and *SF3B1* which were very rare in those homozygous for the alt-allele, suggesting that activation of *TCL1A* expression may be a near requirement for clonal expansion due to these mutations. We do not explore how splicing factor mutations are mechanistically linked to *TCL1A* activation in this study, but one possibility is that mutations in *SF3B1* or *SRSF2* lead to a cryptic splice junction^39,40^ within the *TCL1A* 3’ UTR, which may lead to increased stability of the transcript in HSCs^41^. These results may also potentially explain why mutations in *Asxl1, Sf3b1*, and *Srsf2* in mouse HSCs do not lead to robust clonal expansion, as the regulatory elements and non-coding regions of the mouse *Tcl1* gene are not well conserved with human *TCL1A*. We previously reported that the alt-allele of rs2887399 was associated with increased risk of *DNMT3A* mutations^11^, but here we found that carrying the alt-allele did not increase expansion rate of *DNMT3A* clones by PACER or result in increased expansion of *DNMT3A* edited HSPCs in vitro. One explanation for these discordant results is that the relative reduction in fitness advantage for other non-*DNMT3A* drivers in carriers of the alt-allele permits more opportunity for low fitness *DNMT3A* mutant clones to expand as hematopoiesis becomes more oligoclonal with aging^14^. Alternatively, the interaction of *DNMT3A* mutations with *TCL1A* genotype may not be apparent in middle-aged or older persons, as it has recently been shown that the fitness of *DNMT3A* mutant clones declines with age^17^.

Though our approach yielded several novel insights into clonal expansion, our study has limitations. The sequencing coverage in TOPMed WGS was 38x, which does not enable detection of mutations with VAFs below 5%^11^. More sensitive assays would increase detection of both driver and passenger mutations and would reduce error in estimation of clonal expansion using PACER. Additionally, the use of bulk-WGS precludes analysis of the clonal expansion rate in those with more than one driver mutation. Lastly, we were underpowered to identify germline determinants of clonal expansion in less frequently mutated genes in CHIP, but the addition of WGS data from other cohorts in the coming years should enable additional gene-level analyses of PACER.

In summary, we developed a novel tool for inferring clonal expansion rate and used it to identify *TCL1A* as a factor underlying the clonal fitness advantage of several driver mutations in CHIP. PACER is a powerful approach for identifying the genetic and environmental factors mediating clonal expansion in humans at population scale and may be applied to any tissue where pre-malignant clones exist ^42–44^.

## METHODS

### Study Samples

Whole genome sequencing (WGS) was performed on 127,946 samples as part of 51 studies contributing to Freeze 8 NHLBI TOPMed program as previously described^10^. None of the TOPMed studies included selected individuals for sequencing because of hematologic malignancy. Each of the included studies provided informed consent. Age was obtained for 82,807 of the samples, and the median age was 55, the mean age 52.5, and the maximum age 98. The samples have diverse reported ethnicity (40% European, 32% African, 16% Hispanic/Latino, 10% Asian).

### WGS Processing, Variant Calling and CHIP annotation

BAM files were remapped and harmonized through the functionally equivalent pipeline^45^. SNPs and indels were discovered across TOPMed and were jointly genotyped across samples using the GotCloud pipeline^46^. An SVM filter was trained to discriminate between high- and low-quality variants. Variants were annotated with snpEff 4.3^47^. Sample quality was assessed through mendelian discordance, contamination estimates, sequencing converge, and among other quality control metrics.

Putative somatic SNPs were called with GATK Mutect2^12^, which searches for sites where there is evidence for alt-reads that support evidence for variation, and then performs local haplotype assembly. We used a panel of normals to filter sequencing artifacts and used an external reference of germline variants to exclude germline calls. We deployed this pipeline on Google Cloud using Cromwell^48^.

As described in our previous report ^11^, samples were annotated as having CHIP if the Mutect2 output contained at least one variant in a curated list of leukemogenic driver mutations with at least three alt-reads supporting the call. We expanded the list of driver mutations to include those in recently identified CHIP genes^49^, increasing the number of CHIP cases from our previous report.

We called somatic singletons by identifying somatic variants that appeared in a single individual among the CHIP carriers and 23,320 additional controls for a total of 28,391 individuals. We excluded any variant that appeared in the TOPMed Freeze 5 germline call set (463 million variants). We excluded variants with a depth below 25 or above 100 and excluded any variants in low complexity regions or segmental duplications, as these are challenging for variant calling. We only included somatic singletons that were aligned to the primary chromosomal contigs. We excluded any variant with a VAF exceeding 35% as these may be enriched for germline variants that were not included in our other filters. We used cyvcf2^39^ to parse the Mutect2 VCFs and encoded each variant in an int64 value using the variant key encoding^40^. We developed a bespoke Python application to perform the singleton identification and filtering.

A special approach was required to identify somatic variants in *U2AF1* since an erroneous segmental duplication in the region of the gene in the hg38 reference genome resulted in a mapping score of zero during alignment of the FASTQ file^50^. We developed a Rust-HTSLIB binary (https://github.com/weinstockj/pileup_region) to specifically identify reads associated with the *U2AF1* variants S34F, S34Y, R156H, Q157P, and Q157R. A minimum of 5 alternate reads was required to include a variant in the somatic set of CHIP calls. The variant set was judged to have a high likelihood of being somatic based on the strong age association for persons carrying mutations as well as a high rate of co-mutation with other known drivers. The VAF was estimated by dividing the alternate read count by the total read count for *U2AF1*.

### Amplicon sequencing validation

Targeted sequencing of the CHIP driver genes from 80 samples from the Women’s Health Initiative (WHI) was performed using single-molecule molecular inversion probe sequencing (smMIPS^11,51^). Reads were aligned with bwa-mem and processed with the mimips pileline ^52^. We called somatic variants using an ensemble of VarScan^53^, Mutect2^12^, and manual inspection with IGV^54^.

### Single Variant Association

Single variant association for each variant in the TOPMed Freeze 8 germline genetic variant call set ^10^ with a MAC > 20 was performed with SAIGE^23^ using the TOPMed Encore analysis server. To identify associations between rs2887399 and the acquisition of specific CHIP mutations, we used the same methods as our previous report on an analysis set of 74,974 individuals, including 4,697 cases and 70,277 controls. Age, genotype inferred sex, the first ten genetic ancestry principal components, and study were included as covariates.

We performed SAIGE single variant association analyses on the passengers including age at blood draw, sex, VAF, study, and the first ten genetic ancestry principal components as covariates. We applied an inverse normal transformation to the passenger counts. We declared variants from this analysis as significant if their p-value was less than 5 × 10^−8^.

### Estimation of association between rs2887399 genotypes and CHIP mutation acquisition

We coded the rs2887399 genotypes as a categorical variable rather than a linear quantitative coding to estimate effects separately for the heterozygotes and the alt-homozygotes using the ref-homozygotes as the reference level. We estimated the associations using firth logistic regression to reduce bias in estimation resulting from low cell counts^55^, and included age, genotype inferred sex, and the first ten genetic ancestry components as covariates.

### Fine-mapping of the *TCL1A* region

We applied the SuSIE^24^ algorithm to the genotypes included in a 200kb region surrounding *TCL1A*. We used the same covariates as the single variant association analysis. We used the posterior inclusion probabilities (PIP) and credible sets identified by SuSIE to identify the putative causal variant. We used LD directly calculated on the genotypes as opposed to an external reference.

### Rare Variant Analyses

We performed gene-based tests on 1,698 cancer associated genes their flanking regions using the SCANG^56^ procedure. We identified these genes by downloading the targets associated with cancer in Open Targets^21^, and then filtered to include only genes with an association score of 1.0. The most prevalent CHIP driver genes were included among this list. We used the inverse normal transformed passenger counts as the phenotype with the same covariates as before. We specified the minimum size of the grouped regions as 30 variants and the maximum as 200. We included all PASS variants with a minor allele count greater than four and less than 300 (MAF of 3.7% in the analyzed samples). We parsed the genotypes using cyvcf2^57^ and stored them as dgCMatrix using the Matrix^58^ package from the R 3.6.1 programming language^59^.

We set the p-value filter to calculate SKAT test-statistics at 5 × 10^−4^. We did not group the variants by annotation and we declared regions as significant if their pvalue was less than 2.9 ×10^−5^ (.05 / 1,698). We controlled for relatedness by incorporating a sparse kinship matrix as estimated by the PC-AiR method from the GENESIS R package^60^. We specified separate residual variance terms for each study to control for heterogeneous residual variance. We grouped together all studies where the number of analyzed samples was less than 200.

### Enrichment of passengers by driver gene

We estimated the association between the driver genes and the passenger counts using DNMT3A as the reference in a negative binomial regression using the glm.nb function from the MASS R package^61^. We included age, study, VAF, and sex as covariates. We included driver genes with at least 30 mutations and reported genes that had a different effect relative effect than DNMT3A if the pvalue of the coefficient was less than 1 × 10^−2^.

### Estimation of passenger mutation rate, clone fitness, and clone birth date

We developed a hierarchical Bayesian latent variable model using the Stan^62,63^ probabilistic programming language. We used the negative binomial likelihood with a mean and overdispersion parameterization to facilitate interpretation. We used the identity function to link the passenger counts to the predictors as we modeled the effects on an additive scale. We modeled the expectation and overdispersion of the passenger counts observed at time (*t*_*i*_) as

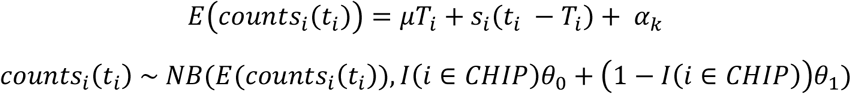

Where *T*_*i*_ is the time of the driver acquisition for sample *i* with a blood draw at time *t*_*i*_, *µ* is the mutation rate per diploid genome per year for the HSC population, *s*_*i*_ is the fitness of the clone, and *a*_*k*_ represents a study specific random intercept for sample *i* included in study *k*. We can interpret *t*_*i*_ − *T*_*i*_ as the lifetime of the clone in years. We used a negative binomial likelihood as there was overdispersion relative to a Poisson distribution.

We included several constraints and priors on the parameters to make them identifiable. We constrained *T*_*i*_ to be positive but exceeded by *t*_*i*_ such that the parameter would be in yearly units. We included case-control specific overdispersion terms *θ*_0_ and *θ*_1_ as the CHIP carriers had greater dispersion. To adjust for batch effects, we included a random intercept, as the amount of singletons in controls varied by study.

To include the constraint on *T*_*i*_, we defined *T*_*i*_ = *ψ*_*i*_ * *age*_*i*_, with *ψ*_*i*_ constrained between 0 and 1, and *age*_*i*_ is the age at blood draw. We placed an uninformative Beta(1, 1.3) prior on *ψ*_*i*_, which is equivalent to the supposition that the driver mutation is twice as likely to be acquired in the second half of life (at the time of blood draw) then the first. We assumed the study specific deviations were exchangeable with respect to a *N*(0,20) prior, providing some shrinkage on the study specific intercepts. We placed a *N*(0,1) prior on the *s*_*i*_ parameter to aid identification. Further details are described in the supplement.

To estimate the posterior, we used the Stan Hamiltonian monte-carlo (HMC) sampler with four separate chains, and used 400 samples of burn-in. We assessed convergence using the Rhat and effect-sample size statistics. We tried multiple parameterizations to reduce the number of divergent transitions. We performed posterior predictive checks to assess the model fit.

### Simulation of HSC dynamics

We simulated the number of cells within an HSC clone as a birth-death continuous time Markov chain, which models the size of an HSC clone as the composite of simultaneous Poisson birth and Poisson death point processes (Supplementary Text 1). Following Watson et al.^20^, HSCs could transition to one of three states: asymmetric renewal, symmetric self-renewal, and symmetric differentiation. The rate of transition was determined by the symmetric differentiation rate of the cell per year, which was set to five. The symmetric self-renewal and symmetric differentiation increase and decrease the size of the HSC clone respectively. As asymmetric division does not affect the size of the clone, we did not explicitly simulate transition to this state. The proclivity towards self-renewal was determined by the fitness of the clone. We set the entire HSC population to acquire a single driver mutation during the ‘lifetime’ of the simulation.

Passengers were accumulated over time using a birth Poisson point process. We then calculated the number of ‘detectable’ passengers that preceded the acquisition of the driver based on whether the underlying clone had expanded to a great enough proportion of HSC cells. We examined the association between the number of detectable passengers and the fitness of the underlying HSC clone. We implemented this simulation in the Julia programming language 1.4^64^.

### Re-analysis of single-cell RNA sequencing data

The cell-by-gene count matrix data for each sample from Psaila et al^36^, generated using the 10X Genomics platform, was downloaded from Gene Expression Omnibus (GSE144568). Each matrix was loaded in Seurat with the read10X command, and only cells with a minimum of 200 features were retained using the CreateSeuratObject command. Data was log normalized using a scale factor of 10000 by the NormalizeData command. We then used the FindVariableFeatures command with ‘vst’ selection method and 2000 features. The data was scaled using ScaleData using all genes as features. We then used the RunPCA command with VariableFeatures identified earlier. For clustering, we used FindNeighbors set to the first 10 PCA dimensions and FindClusters using a resolution of 0.5. We excluded samples that did not have a distinct cluster of HSC/MPPs, defined as clusters enriched for cells that were *CD34*+ *CD38*-/lo *THY1*+. This left 5 healthy marrow samples (id01, id06, id09, id13, id17) and 4 MPN samples (id2, id7, id11, id14). For each of these samples, we assessed the number of cells with *TCL1A* transcripts within the cluster or clusters that contained HSC/MPPs, as defined above.

Additional preprocessed single-cell RNAseq from Velten et al.^35^, generated using MutaSeq, was downloaded from Gene Expression Omnibus (GSE75478) as an RDS file. We utilized data from one patient with AML (P1) and the healthy control (H1). We then determined the number of cells containing *TCL1A* transcript in the preleukemic ‘HSC/MPP’ and preleukemic ‘CD34+ blasts and HSPCs’ clusters for the P1 sample and the ‘HSC/MPP’ cluster for the H1 sample, in both cases as defined by the original study authors.

### Re-analysis of ATACseq data

We downloaded ATAC-seq data for AML samples as well as healthy controls from Corces et al.^65^ available at Gene Expression Omnibus (GSE75478). For our analysis, we used data from HSCs, defined as Lin-CD34+ CD38-CD90+ CD10-by the authors, from a healthy donor (donor7256), or preleukemic HSCs (pHSC), defined as Lin-CD34+ CD38-TIM3-CD99-by the authors. For the pHSC samples, we selected 2 where there were no detectable driver mutations in the pHSC compartment (SU336, SU306), 2 where there were *DNMT3A* mutations only (SU444, SU575), and 2 where there were *TET2* mutations only (SU070, SU501).

Fastq files were downloaded, and ATAC-seq data analysis was performed as previously described^66^.

Briefly, reads were trimmed and filtered using fastp and mapped to the hg38 reference genome using hisat2 with the --no-spliced-alignment option. Bam files were deduplicated using Picard. Only reads mapping to chromosomes 1-22 and chrX were retained -- chrY reads, mitochondrial reads, and other reads were discarded. Genome track files were created by loading the fragments for each sample into R, and exporting bigwig files normalized by reads in transcription start sites using ‘rtracklayer::export’. Coverage files were visualized using the Integrative Genomics Viewer.

### CRISPR–Cas9 editing of CD34^+^ human HSPCs

CD34^+^ HSPCs from adult donors were purchased from the Cooperative Center of Excellence in Hematology (CCEH) at the Fred Hutch Cancer Research Center, Seattle, USA. TCL1A rs2887399 genotyping was performed using ThermoFisher SNP assay (Assay ID: C 15842295_20). CD34+ cells were thawed and cultured in HSC Expansion media (StemSpanII + 10% CD34+ Expansion Supplement + 0.1% Penicillin/Streptomycin) for 48 hours before CRISPR editing. Editing of *AAVS, TET2, DNMT3A*, and *ASXL1* was performed by electroporation of Cas9 ribonucleoprotein complex (RNP). For each combination of rs2887399 genotype and gRNA, 100,000 cells were incubated with 3.2 ug of Synthego synthetic sgRNA guide and 8.18 ug of IDT Alt-R S.p. Cas9 Nuclease V3 for 15 minutes at room temperature before electroporation. CD34+ cells were resuspended in 18 uL of Lonza P3 solution and mixed with the ribonucleoprotein complex, and then transferred to Nucleocuvette strips for electroporation with program DZ-100 (Lonza 4D Nucleofector). Immediately following electroporation, each condition of 100,000 cells was transferred to 2 mL’s of HSC Expansion media, and allowed to recover for 24 hours. CRISPR editing efficiency was measured using Sanger Sequencing and ICE Analysis.

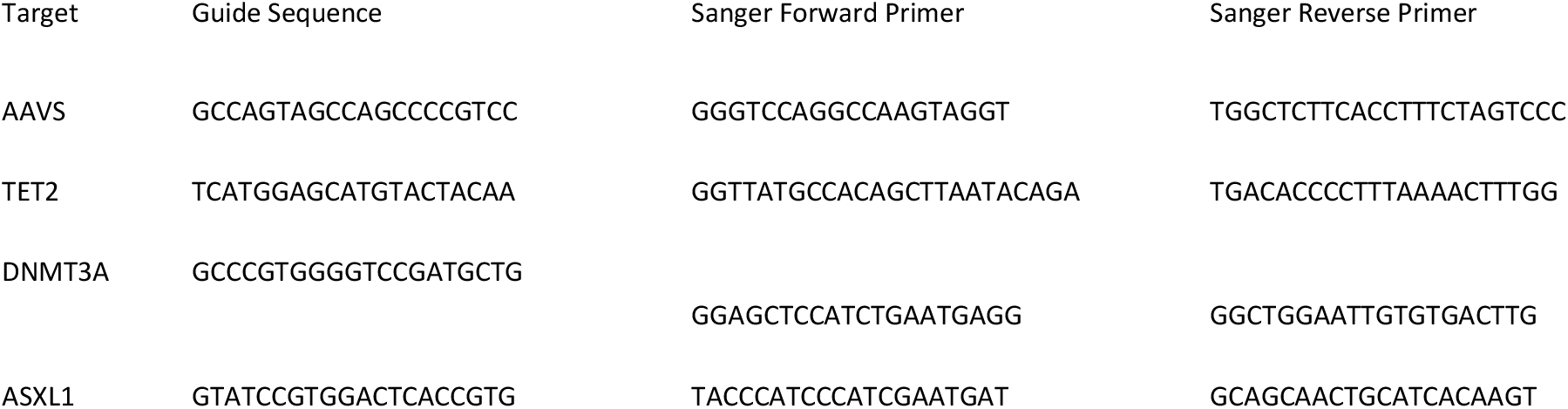

### Liquid Culture Expansion Assay

24 hours post electroporation, Lineage-CD34+ CD38-CD90+ CD45RA-cells were sorted on a BD FACS Aria III from the electroporated CD34+ cells All cells were harvested and stained with the following extracellular HSC marker panel in 100 uL of PBS + 2% FBS + 1 mm EDTA.

| Antibody | Vendor | Clone | Catalog# | Concentration |
| --- | --- | --- | --- | --- |
| APC-CD34 | BioLegend | 561 | 343608 | 1:50 |
| PE/Cy7-CD38 | BioLegend | HIT2 | 303515 | 1:100 |
| BV421-CD90 | BioLegend | 5E10 | 562556 | 1:25 |
| BV605-CD45RA | BioLegend | HI100 | 304134 | 1:40 |
| PE/Cy5-CD2 | BioLegend | RPA-2.10 | 300210 | 1:100 |
| PE/Cy5-CD3 | BioLegend | HIT3a | 300310 | 1:100 |
| PE/Cy5-CD4 | BioLegend | SK3 | 344654 | 1:100 |
| PE/Cy5-CD8a | BioLegend | HIT8a | 300910 | 1:100 |
| PE/Cy5-CD16 | BioLegend | 3G8 | 302010 | 1:100 |
| PE/Cy5-CD19 | BioLegend | HIB19 | 302210 | 1:100 |
| PE/Cy5-CD20 | BioLegend | 2H7 | 302308 | 1:100 |
| PE/Cy5-CD56 | BioLegend | 5.1H11 | 362516 | 1:100 |
| PE/Cy5-CD235a | BioLegend | HIR2 | 306606 | 1:100 |
| PE/Cy5-CD14 | BioLegend | M5E2 | 301864 | 1:100 |
| Fixable Viability Stain 700 | BD |  | 564997 | 1:1000 |

4 replicates of 1,000 Lineage-CD34+ CD38-CD90+ CD45RA-cells were sorted into 100 uL of HSC Expansion media and cells were plated into a 96 well plate. The edges of 96 well plate were filled with water to keep the cultures hydrated. 4 days post sort, another 100 uL of HSC Expansion media was added to each well. 10 days post sort, the samples were transferred from the 96 well plate to a 48 well plate and an additional 400 uL of HSC Expansion media was added. 14 days post sort, the cells were harvested, and live cells were counted using trypan blue and hemocytometer. Additionally, the cells were stained with the extracellular HSC marker panel, and flow cytometry analysis was performed. Absolute number of HSC/MPPs (defined as Lin-CD34+ CD38-CD45RA-) and CD45RA^lo^progenitors (defined as Lin-/lo CD34+ CD38-CD45RA^lo^) were determined by multiplying the total cell count at 14 days by the percentage of cells in each compartment as determined by flow cytometry.

### Flow cytometry for TCL1A staining

Anti-human TCL1A antibody clone eBio1-21 was obtained from ThermoFisher. The specificity of the antibody was assessed by staining NALM6 cells that had been CRISPR edited for *TCL1A* with the antibody, which confirmed only a low level of non-specific binding.

To assess for TCL1A expression in cultured human HSPCs, cells in HSC Expansion media were harvested and intracellularly stained 11 days following electroporation.

| Antibody | Vendor | Clone | Catalog# | Concentration |
| --- | --- | --- | --- | --- |
| e450-TCL1A | ThermoFisher | eBio1-21 | 48-6699-42 | 1 ug |
| APC-CD34 | BioLegend | 561 | 343608 | 1:50 |
| PE/Cy7-CD38 | BioLegend | HIT2 | 303515 | 1:100 |
| FITC-CD90 | BioLegend | 5E10 | 562556 | 1:15 |
| BV605-CD45RA | BioLegend | HI100 | 304134 | 1:25 |
| PE/Cy5-CD2 | BioLegend | RPA-2.10 | 300210 | 1:100 |
| PE/Cy5-CD3 | BioLegend | HIT3a | 300310 | 1:100 |
| PE/Cy5-CD4 | BioLegend | SK3 | 344654 | 1:100 |
| PE/Cy5-CD8a | BioLegend | HIT8a | 300910 | 1:100 |
| PE/Cy5-CD16 | BioLegend | 3G8 | 302010 | 1:100 |
| PE/Cy5-CD19 | BioLegend | HIB19 | 302210 | 1:100 |
| PE/Cy5-CD20 | BioLegend | 2H7 | 302308 | 1:100 |
| PE/Cy5-CD56 | BioLegend | 5.1H11 | 362516 | 1:100 |
| PE/Cy5-CD235a | BioLegend | HIR2 | 306606 | 1:100 |
| PE/Cy5-CD14 | BioLegend | M5E2 | 301864 | 1:100 |
| Fixable Viability Stain 700 | BD |  | 564997 | 1:1000 |

Cells were first stained with the Live/Dead and extracellular HSC markers simultaneously for 30 minutes in the dark on ice. After a PBS wash, cells were stained with 100 uL of IC Fixation Buffer for 30 minutes in the dark at room temperature. Cells were then washed twice with 1X Permeabilization Buffer. Next, cells were resuspended in 100 uL of 1X Permeabilization Buffer, and blocked with 2 uL of goat serum and 2.5 uL of TruStain FcX for 15 minutes in the dark at room temperature. Next, 1 ug of e450 antibodies (anti-TCL1A or isotype control) was added to each sample tube and stained for 30 minutes in the dark at room temperature. Cells were then washed twice with 1X Permeabilization Buffer and then resuspended in PBS before flow cytometry was performed. HSC/MPPs were defined as Lin-CD34+ CD38-CD45RA-.

### ATAC-seq

24 hours post electroporation, Lineage-CD34+ CD38-CD45RA-cells were sorted from the electroporated CD34+ cells using a BD FACS Aria III. Cells were allowed to culture for 5 days before 40,000 cells were harvested, and bulk Omni-ATAC^67^ was performed on them. Briefly, cells were lysed with ATAC-Resuspension Buffer containing 0.1% NP40, 0.1% Tween-20, and 0.01% Digitonin for 3 minutes, and then the transposition was performed for 30 minutes at 37 C using 100 nM of Illumina Tagment DNA TDE1 Enzyme and Buffer Kit per 50,000 cells. The fragmented DNA was then cleaned up using a Zymo DNA Clean and Concentrator-5 Kit (cat# D4014). The transposed fragments were amplified and indexed using NEBNext 2x Master Mix. The final PCR product was purified using the Zymo DNA Clean and Concentrator-5 Kit. Prior to sequencing, the quality of the libraries was evaluated via DNA High Sensitivity Bioanalyzer assays. The sequencing was performed using 2×75 bp reads on an Illumina NextSeq550 instrument using the High Output Kit.

ATAC-seq data analysis was performed as previously described above. Briefly, reads were trimmed and filtered using fastp and mapped to the hg38 reference genome using hisat2 with the --no-spliced-alignment option. Bam files were deduplicated using Picard. Only reads mapping to chromosomes 1-22 and chrX were retained -- chrY reads, mitochondrial reads, and other reads were discarded. Genome track files were created by loading the fragments for each sample into R, and exporting bigwig files normalized by reads in transcription start sites using ‘rtracklayer::export’.

Coverage files were visualized using the Integrative Genomics Viewer.

## Supporting information

Supplementary Acknowledgements

Supplementary Theoretical Text

## DATA AVAILABILITY

Individual whole-genome sequence data for TOPMed whole genomes, individual-level harmonized phenotypes and the CHIP variant call sets used in this analysis are available through restricted access via the dbGaP TOPMed Exchange Area available to TOPMed investigators. Controlled-access release to the general scientific community via dbGaP is ongoing.

## CODE AVAILABILITY

https://github.com/weinstockj/hsc_simulation

https://github.com/weinstockj/pileup_region

https://github.com/weizhouUMICH/SAIGE https://github.com/zilinli1988/SCANG

https://dockstore.org/workflows/github.com/broadinstitute/gatk/mutect2:4.1.8.1?tab=info

https://stephenslab.github.io/susieR/index.html

## ACKNOWLEDGEMENTS

WGS for the TOPMed program was supported by the National Heart, Lung and Blood Institute (NHLBI). Centralized read mapping and genotype calling, along with variant quality metrics and filtering were provided by the TOPMed Informatics Research Center (3R01HL-117626-02S1; contract HHSN268201800002I). Phenotype harmonization, data management, sample-identity quality control and general study coordination were provided by the TOPMed Data Coordinating Center (R01HL-120393; U01HL-120393; contract HHSN268201800001I). We thank the studies and participants who provided biological samples and data for TOPMed. The full study-specific acknowledgments are included in Supplementary Information 1. The views expressed in this manuscript are those of the authors and do not necessarily represent the views of the National Heart, Lung, and Blood Institute; the National Institutes of Health; or the US Department of Health and Human Services. The authors wish to acknowledge the contributions of the consortium working on the development of the NHLBI BioData Catalyst ecosystem. S.J. is supported by the Burroughs Wellcome Foundation Career Award for Medical Scientists, Foundation Leducq, Ludwig Center for Cancer Stem Cell Research, the American Society of Hematology Scholar Award, and the NIH Director’s New Innovator Award (DP2-HL157540). A.G.B. is supported by a Burroughs Wellcome Foundation Career Award for Medical Scientists and the NIH Director’s Early Independence Award (DP5-OD029586). We thank Ravi Majeti and Thomas Koehnke for helpful discussions.

## FIGURES

**Extended Data Fig 1.**
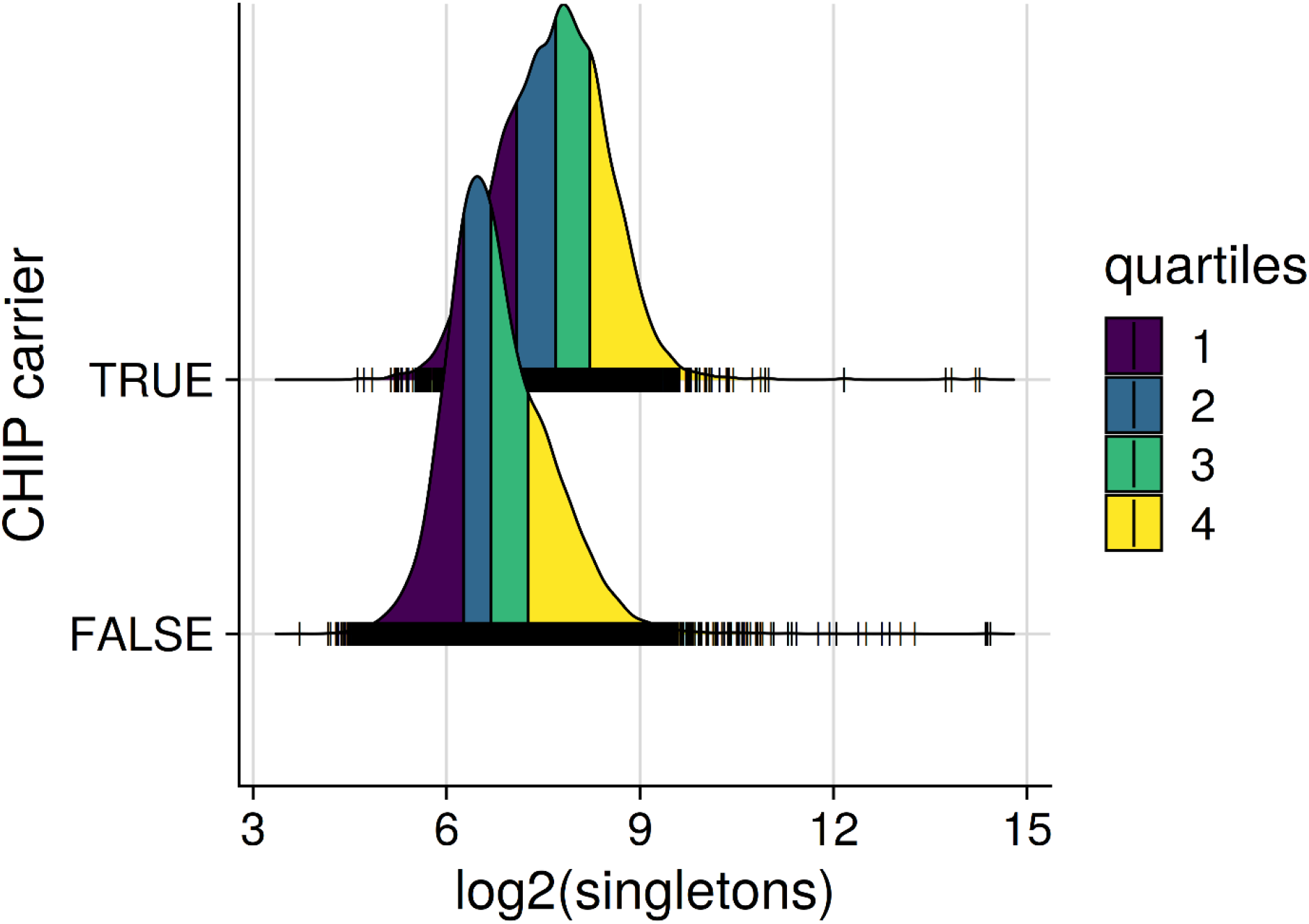
CHIP Carriers are Enriched for Passengers. The passenger counts are enriched by 54% (95% CI: 51%-57%) after adjusting for age and study using a negative binomial regression. The different colors in the density plots correspond to quartiles of the marginal probability distributions. As the density estimates are smoothed, the underlying data points are indicated with hash marks. The data use a log2 scale, such that an increase by 1 indicates a single doubling has occurred.

**Extended Data Fig 2.**
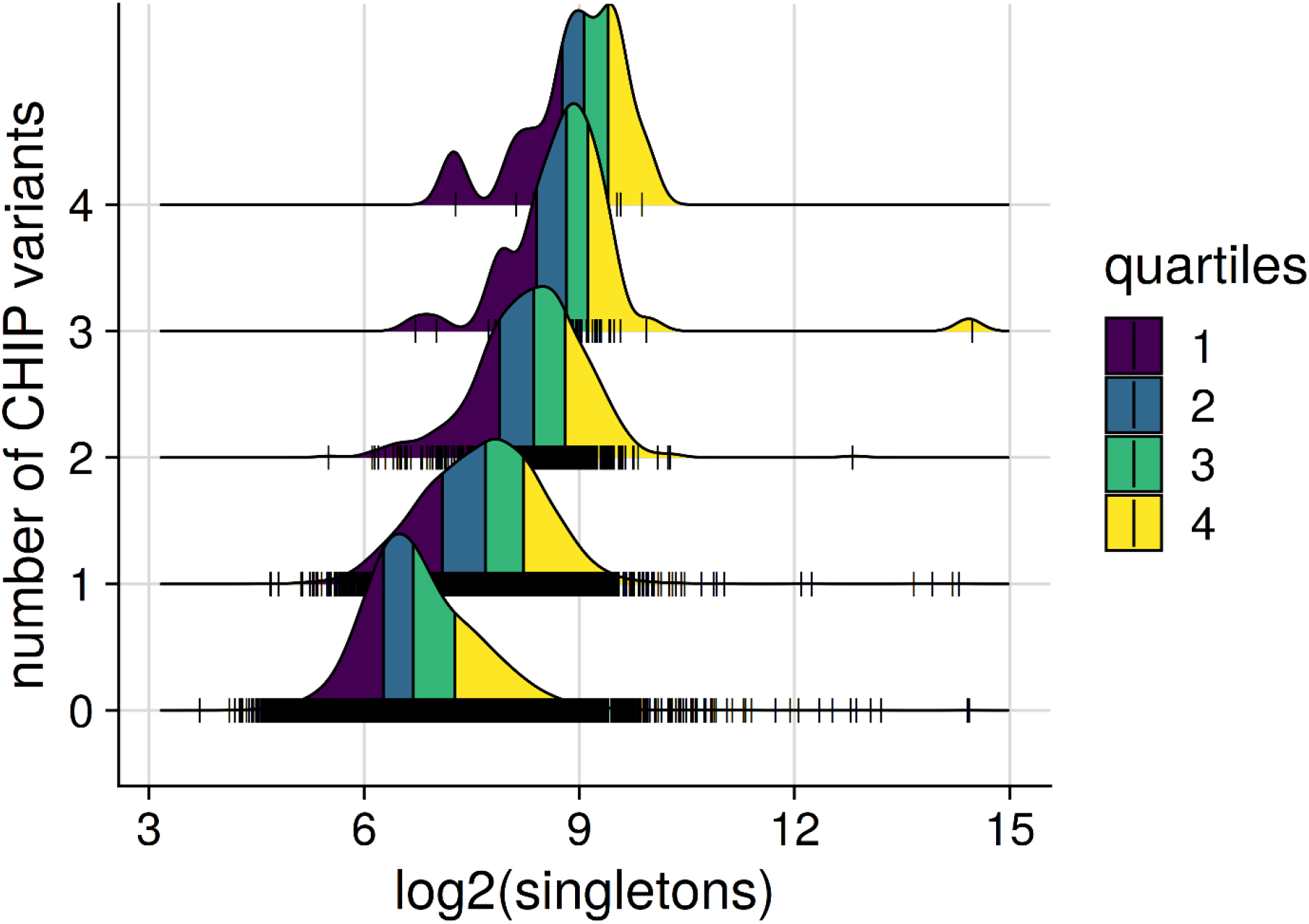
Passenger Counts Linearly Increase with Number of Driver Mutations. The distributions of passenger counts are stratified by the number of CHIP driver variants acquired. The different colors in the density plots correspond to quartiles of the marginal probability distributions.

**Extended Data Fig 3.**
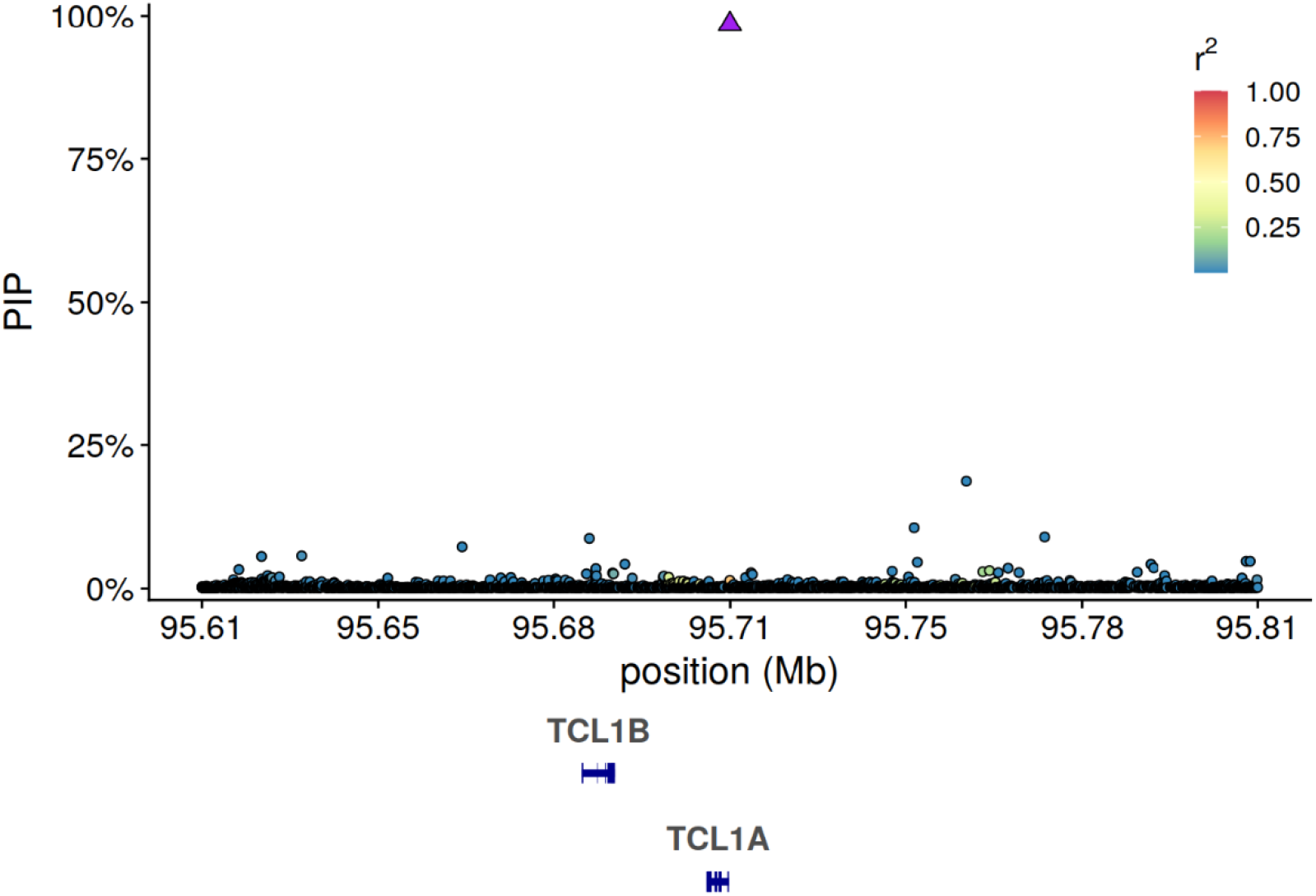
Fine-mapping *TCL1A* Locus Identifies a Single Causal Variant rs2887399. The posterior inclusion probabilities (PIP) as estimated by SuSIE^24^ are plotted on the y-axis, and the genomic position of a 0.8 Mb region including TCL1A is plotted on the x-axis. The linkage disequilibrium (LD) estimates are plotted on a color scale and are estimated on the genotypes used for association analyses.

**Extended Data Fig 4.**
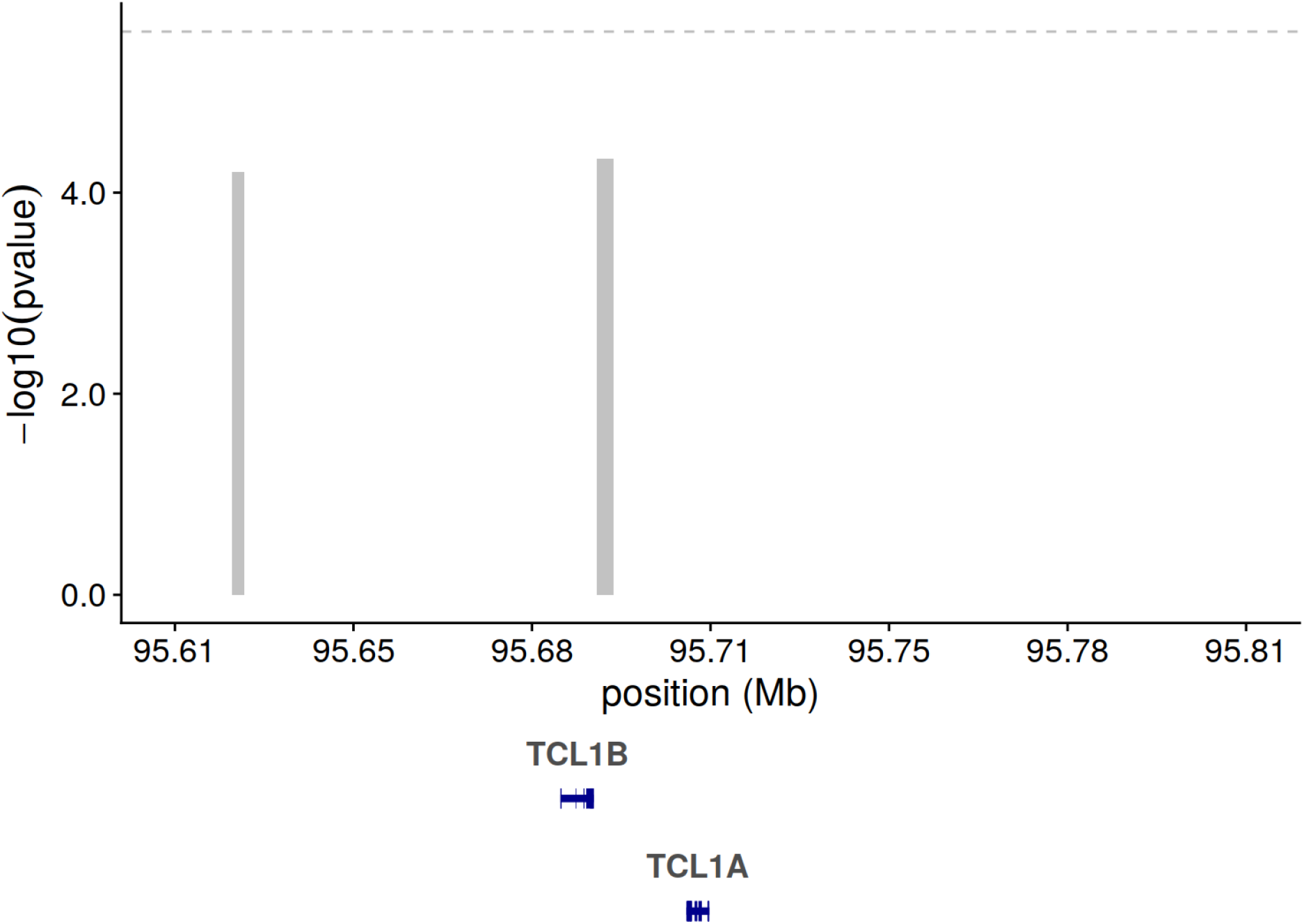
Rare Variant Analysis Of TCL1A Locus Identifies a Suggestive Signal Prior to Conditioning on rs2887399. Rare variant analyses were performed using the SCANG^56^ rare variant scan procedure including all variants with a minor allele count less than 300. Identified rare variant windows are plotted as gray rectangles where the width corresponds to the size of the genomic region and the height corresponds to the pvalue of the SCANG test statistic for the window.

**Extended Data Fig 5.**
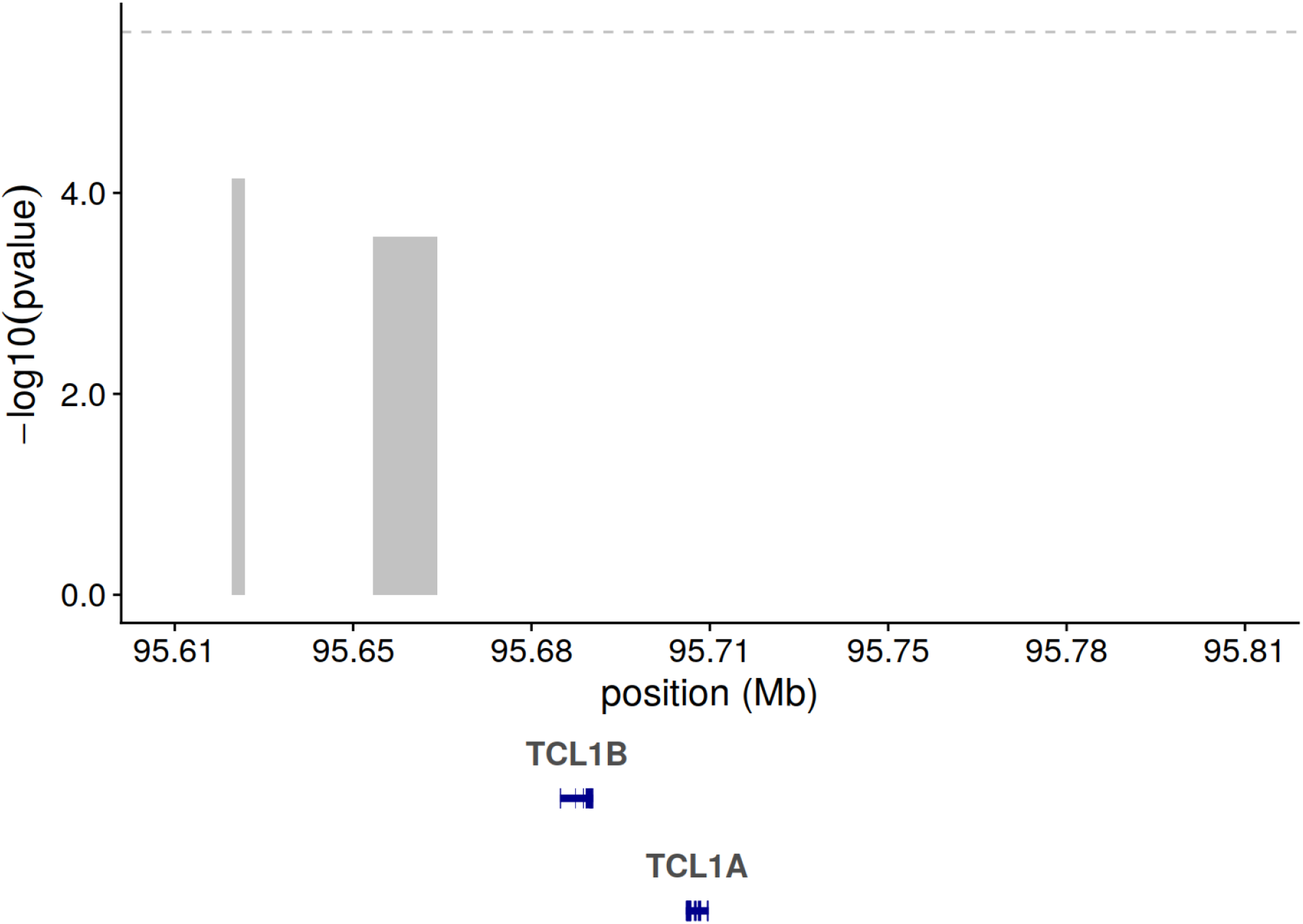
Conditioning on rs2887399 Attenuates Independent Rare Variant Signal. Rare variant analyses were performed including the rs2887399 genotypes as covariate.

**Extended Data Fig 6.**
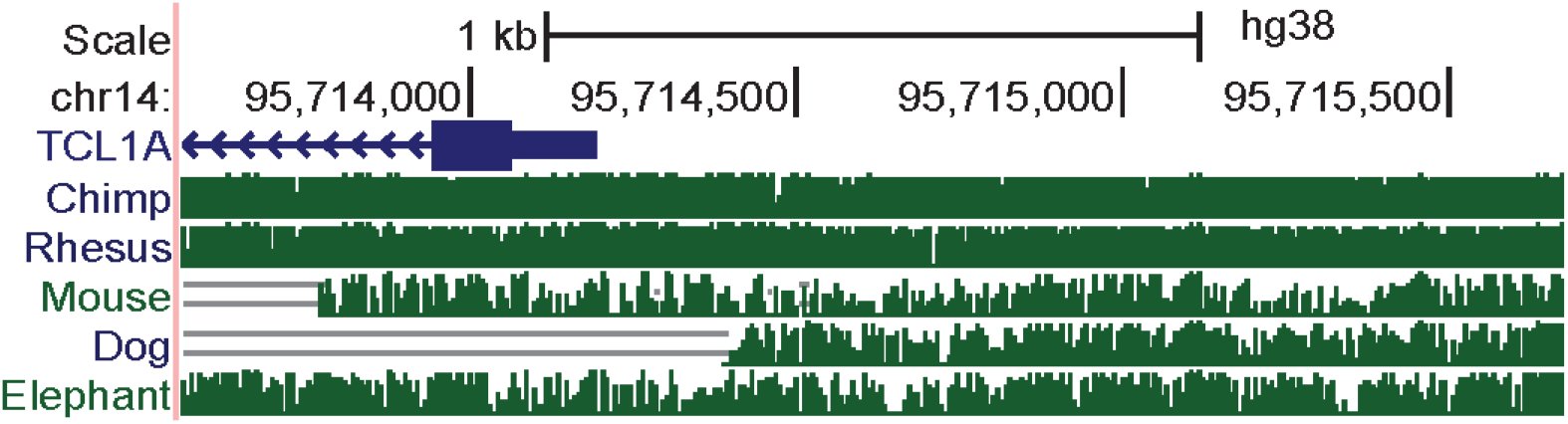
TCL1A Promoter is Not Well Conserved In Vertebrates. Multiz alignments across multiple species are shown for the TCL1A locus.

**Extended Data Fig 7.**
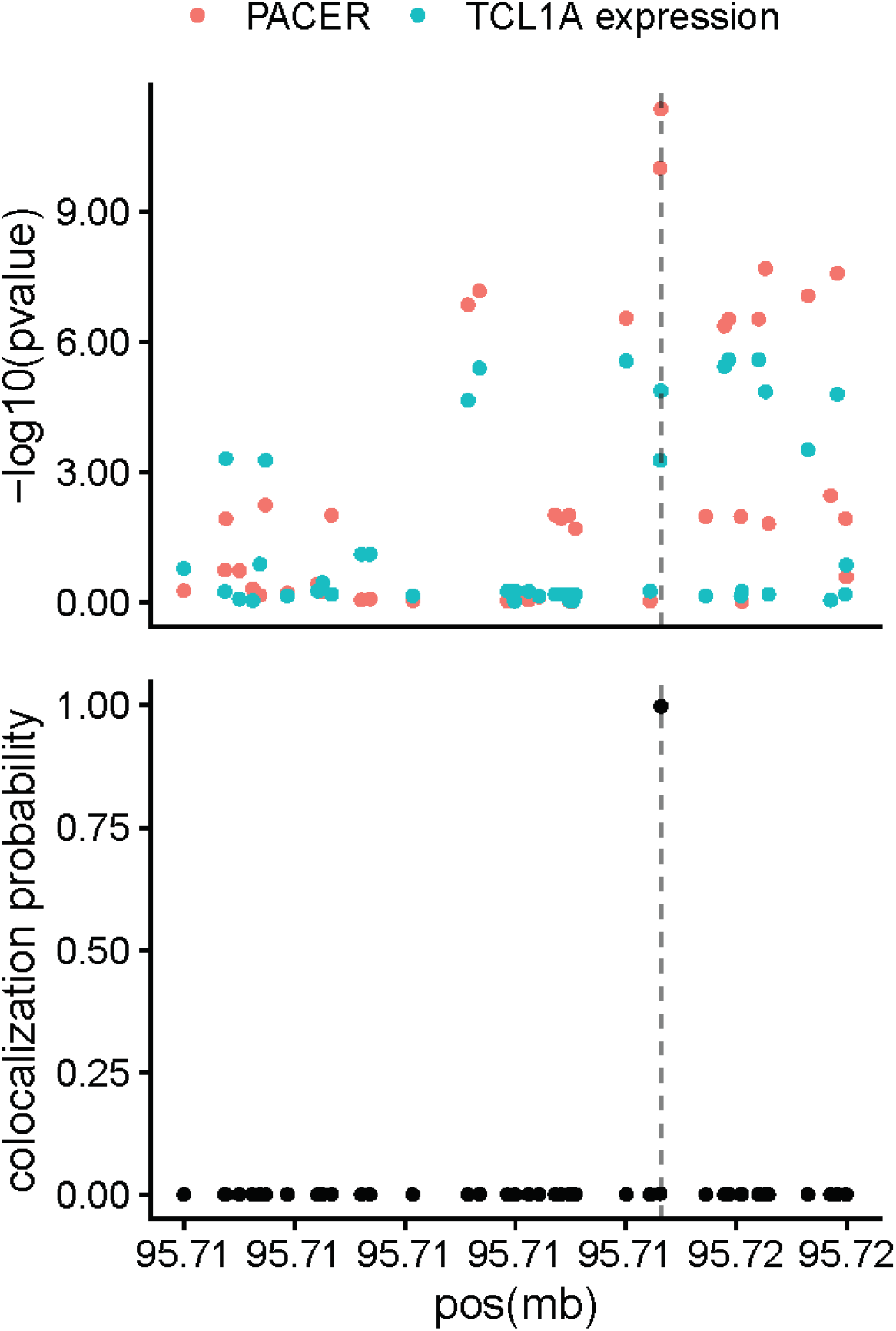
PACER Signal Colocalizes with TCL1A eQTLs. In the top panel, plotted are the -log10 pvalues from both the PACER GWAS and TCL1A cis-eQTLs in whole blood from GTEx v8. In the bottom panel, posterior probability of colocalization from COLOC^68^ identifies rs2887399 as the likely shared causal variant.

**Extended Data Figure 8.**
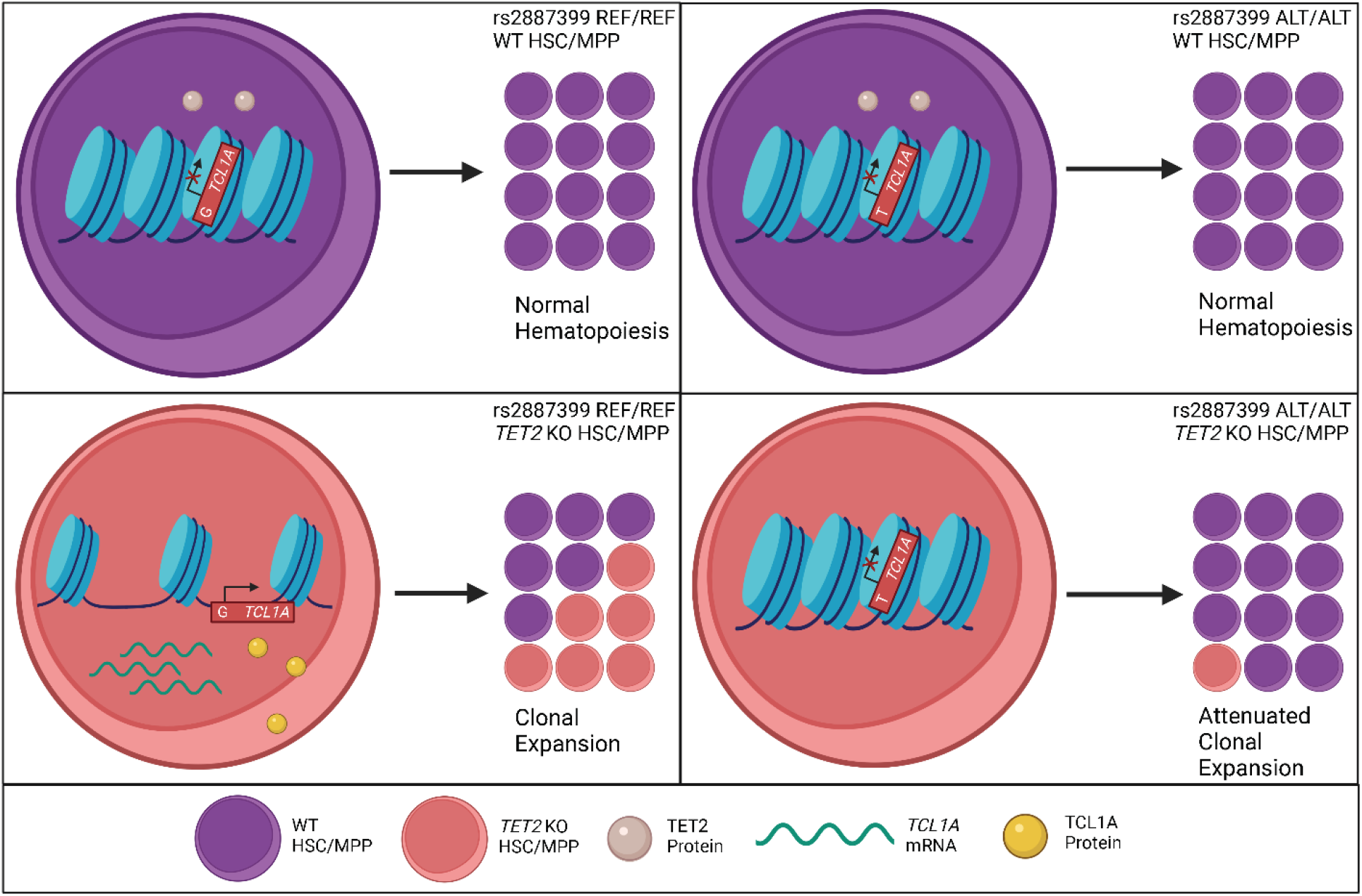
Schematic Description of rs2887399 Mediation on TET2 Clonal Expansion. Proposed model for clonal advantage due to mutations in *TET2*. In cells with the rs2887399 REF/REF genotype, loss of *TET2* function leads to an accessible *TCL1A* locus, aberrant *TCL1A* RNA and protein expression in hematopoietic stem cells (HSC’s) and multi-potent progenitors (MPP’s), and subsequent clonal expansion. The presence of rs2887399 ALT alleles diminishes the *TET2* clonal expansion phenotype by limiting *TCL1A* locus accessibility and downstream protein expression.

**Extended Data Figure 9.**
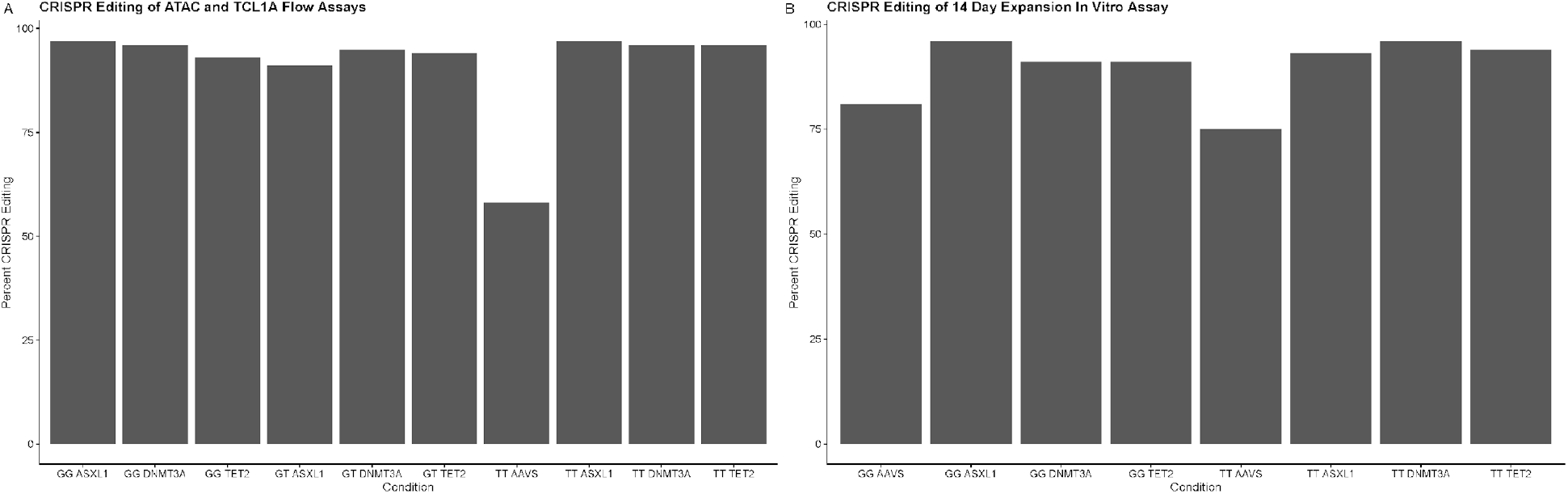
CRISPR Editing Efficiency. **A**. ICE analysis of Sanger traces to determine targeted CRISPR editing efficiency. Bar plots display percent of CD34+ CD38-CD45RA-cells with indel formation in gene of interest. These cells were used for the OMNI-ATAC and intracellular TCL1A flow assays. **B**. ICE analysis of Sanger traces to determine targeted CRISPR editing efficiency. Bar plots display percent of CD34+ CD38-CD45RA-cells with indel formation in gene of interest. These cells were used for the 14 day expansion assay.

**Extended Data Figure 10.**
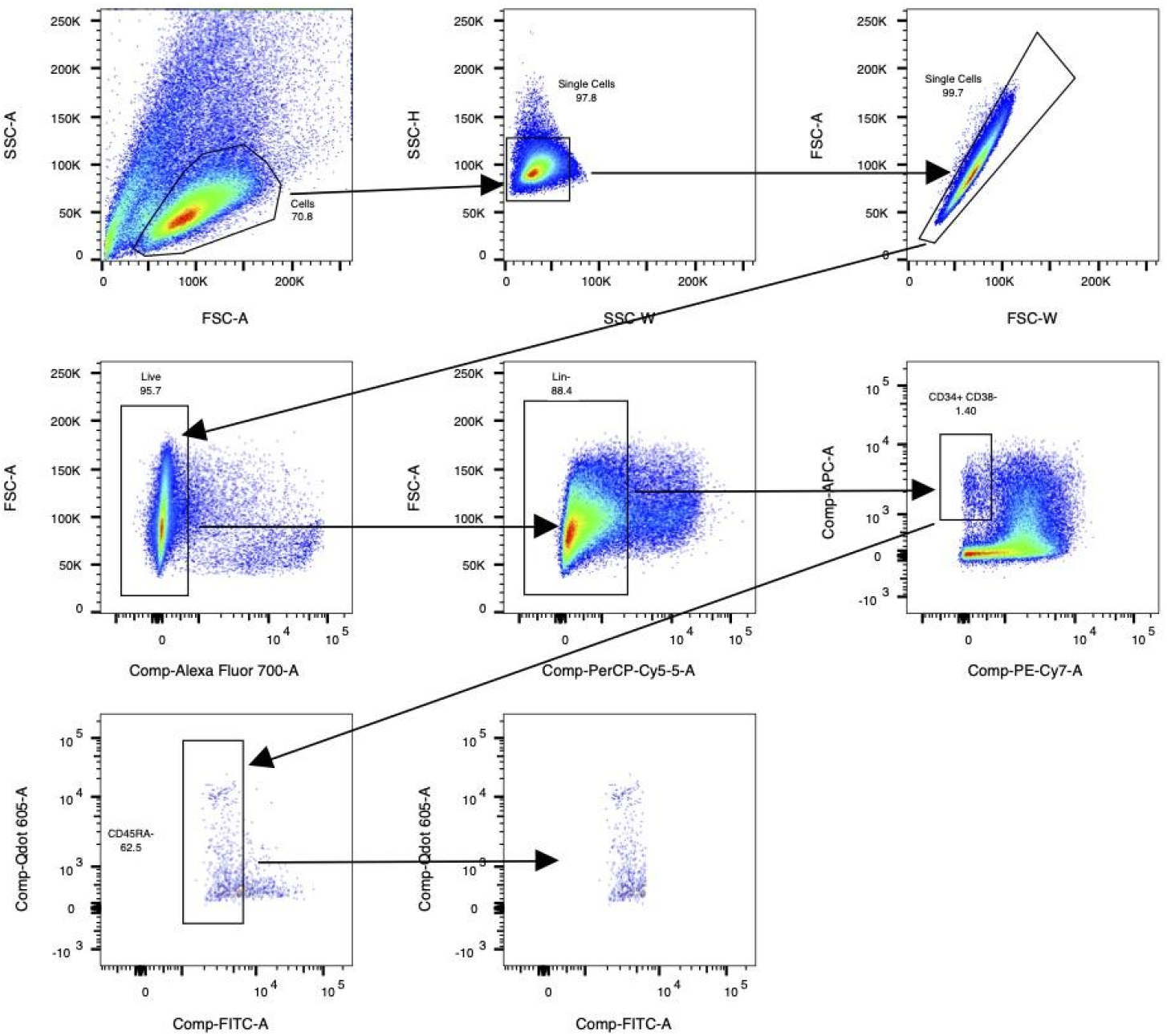
HSC/MPP Flow Gating Scheme. Flow gating scheme for identifying and sorting CD34+ CD38-CD45RA-hematopoietic stem cells (HSC’s) and multi-potent progenitors (MPP’s).

## WORKS CITED

1. Steensma, D. P. et al. Clonal hematopoiesis of indeterminate potential and its distinction from myelodysplastic syndromes. Blood 126, 9–16 (2015).

2. Jaiswal, S. et al. Age-Related Clonal Hematopoiesis Associated with Adverse Outcomes A BS TR AC T. NEJM.org. N Engl J Med 26, 2488–98 (2014).

3. Genovese, G. et al. Clonal Hematopoiesis and Blood-Cancer Risk Inferred from Blood DNA Sequence. New England Journal of Medicine 371, 2477–2487 (2014).

4. Xie, M. et al. Age-related mutations associated with clonal hematopoietic expansion and malignancies. Nature Medicine 20, 1472–1478 (2014).

5. Abelson, S. et al. Prediction of acute myeloid leukaemia risk in healthy individuals. Nature 559, 400–404 (2018).

6. Desai, P. et al. Somatic mutations precede acute myeloid leukemia years before diagnosis. Nature Medicine 24, 1015–1023 (2018).

7. Young, A. L., Challen, G. A., Birmann, B. M. & Druley, T. E. Clonal haematopoiesis harbouring AML-associated mutations is ubiquitous in healthy adults. Nat Commun 7, 12484 (2016).

8. Jaiswal, S. et al. Clonal Hematopoiesis and Risk of Atherosclerotic Cardiovascular Disease. New England Journal of Medicine (2017) doi:10.1056/NEJMoa1701719.

9. Bick Alexander G. et al. Genetic Interleukin 6 Signaling Deficiency Attenuates Cardiovascular Risk in Clonal Hematopoiesis. Circulation 141, 124–131 (2020).

10. Taliun, D. et al. Sequencing of 53,831 diverse genomes from the NHLBI TOPMed Program. Nature 590, 290–299 (2021).

11. Bick, A. G. et al. Inherited causes of clonal haematopoiesis in 97,691 whole genomes. Nature 586, 763–768 (2020).

12. Cibulskis, K. et al. Sensitive detection of somatic point mutations in impure and heterogeneous cancer samples. Nature Biotechnology 31, 213–219 (2013).

13. Osorio, F. G. et al. Somatic Mutations Reveal Lineage Relationships and Age-Related Mutagenesis in Human Hematopoiesis. Cell Reports 25, 2308-2316.e4 (2018).

14. Mitchell, E. et al. Clonal dynamics of haematopoiesis across the human lifespan. 2021.08.16.456475 https://www.biorxiv.org/content/10.1101/2021.08.16.456475v1 (2021) doi:10.1101/2021.08.16.456475.

15. Williams, N. et al. Phylogenetic reconstruction of myeloproliferative neoplasm reveals very early origins and lifelong evolution. 2020.11.09.374710 https://www.biorxiv.org/content/10.1101/2020.11.09.374710v1 (2020) doi:10.1101/2020.11.09.374710.

16. Lee-Six, H. et al. Population dynamics of normal human blood inferred from somatic mutations. Nature 561, 473–478 (2018).

17. Fabre, M. A. et al. The longitudinal dynamics and natural history of clonal haematopoiesis. 2021.08.12.455048 https://www.biorxiv.org/content/10.1101/2021.08.12.455048v1 (2021) doi:10.1101/2021.08.12.455048.

18. Alexandrov, L. B. et al. Clock-like mutational processes in human somatic cells. Nature Genetics 47, 1402–1407 (2015).

19. Zink, F. et al. Clonal hematopoiesis, with and without candidate driver mutations, is common in the elderly. Blood 130, 742–752 (2017).

20. Watson, C. J. et al. The evolutionary dynamics and fitness landscape of clonal hematopoiesis. Science 367, 1449–1454 (2020).

21. Deuren, R. C. van et al. Clone expansion of mutation-driven clonal hematopoiesis is associated with aging and metabolic dysfunction in individuals with obesity. 2021.05.12.443095 https://www.biorxiv.org/content/10.1101/2021.05.12.443095v1 (2021) doi:10.1101/2021.05.12.443095.

22. Robertson, N. A. et al. Longitudinal dynamics of clonal hematopoiesis identifies gene-specific fitness effects. 2021.05.27.446006 https://www.biorxiv.org/content/10.1101/2021.05.27.446006v3 (2021) doi:10.1101/2021.05.27.446006.

23. Zhou, W. et al. Efficiently controlling for case-control imbalance and sample relatedness in large-scale genetic association studies. Nature Genetics 50, 1335–1341 (2018).

24. Wang, G., Sarkar, A., Carbonetto, P. & Stephens, M. A simple new approach to variable selection in regression, with application to genetic fine mapping. Journal of the Royal Statistical Society: Series B (Statistical Methodology) 82, 1273–1300 (2020).

25. Dr, Z., Sp, W., n, J., t, J. & Pr, F. The ensembl regulatory build. Genome Biol 16, 56–56 (2015).

26. Carvalho-Silva, D. et al. Open Targets Platform: new developments and updates two years on. Nucleic Acids Res 47, D1056–D1065 (2019).

27. Narducci, M. G. et al. TCL1 Is Overexpressed in Patients Affected by Adult T-Cell Leukemias. Cancer Res 57, 5452–5456 (1997).

28. Saliba, J. et al. Germline duplication of ATG2B and GSKIP predisposes to familial myeloid malignancies. Nat Genet 47, 1131–1140 (2015).

29. Babushok, D. V. et al. Germline duplication of ATG2B and GSKIP genes is not required for the familial myeloid malignancy syndrome associated with the duplication of chromosome 14q32. Leukemia 32, 2720–2723 (2018).

30. Fishilevich, S. et al. GeneHancer: genome-wide integration of enhancers and target genes in GeneCards. Database (Oxford) 2017, (2017).

31. Thompson, D. J. et al. Genetic predisposition to mosaic Y chromosome loss in blood. Nature 575, 652–657 (2019).

32. Malcovati, L. et al. Clinical significance of somatic mutation in unexplained blood cytopenia. Blood 129, 3371–3378 (2017).

33. Bycroft, C. et al. Genome-wide genetic data on ∼500,000 UK Biobank participants. bioRxiv 166298–166298 (2017) doi:10.1101/166298.

34. Consortium, T. Gte. The GTEx Consortium atlas of genetic regulatory effects across human tissues. Science 369, 1318–1330 (2020).

35. Velten, L. et al. Identification of leukemic and pre-leukemic stem cells by clonal tracking from single-cell transcriptomics. Nat Commun 12, 1366 (2021).

36. Psaila, B. et al. Single-Cell Analyses Reveal Megakaryocyte-Biased Hematopoiesis in Myelofibrosis and Identify Mutant Clone-Specific Targets. Mol Cell 78, 477-492.e8 (2020).

37. Zhou, W. et al. Mosaic loss of chromosome Y is associated with common variation near TCL1A. Nature Genetics 48, 563–568 (2016).

38. Laine, J., Künstle, G., Obata, T., Sha, M. & Noguchi, M. The Protooncogene TCL1 Is an Akt Kinase Coactivator. Molecular Cell 6, 395–407 (2000).

39. Lee, S. C.-W. et al. Synthetic Lethal and Convergent Biological Effects of Cancer-Associated Spliceosomal Gene Mutations. Cancer Cell 34, 225-241.e8 (2018).

40. Obeng, E. A. et al. Physiologic expression of SF3B1K700E causes impaired erythropoiesis, aberrant splicing, and sensitivity to pharmacologic spliceosome modulation. Cancer Cell 30, 404–417 (2016).

41. Mayr, C. What Are 3′ UTRs Doingã Cold Spring Harb Perspect Biol 11, a034728 (2019).

42. Martincorena, I. et al. High burden and pervasive positive selection of somatic mutations in normal human skin. Science 348, 880–886 (2015).

43. Kakiuchi, N. & Ogawa, S. Clonal expansion in non-cancer tissues. Nature Reviews Cancer 21, 239–256 (2021).

44. Martincorena, I. et al. Somatic mutant clones colonize the human esophagus with age. Science 362, 911–917 (2018).

45. Regier, A. A. et al. Functional equivalence of genome sequencing analysis pipelines enables harmonized variant calling across human genetics projects. Nature Communications 9, 1–8 (2018).

46. Jun, G., Wing, M. K., Abecasis, G. R. & Kang, H. M. An efficient and scalable analysis framework for variant extraction and refinement from population scale DNA sequence data. Genome Res. gr.176552.114 (2015) doi:10.1101/gr.176552.114.

47. Cingolani, P. et al. A program for annotating and predicting the effects of single nucleotide polymorphisms, SnpEff: SNPs in the genome of Drosophila melanogaster strain w1118; iso-2; iso-3. Fly 6, 80–92 (2012).

48. Voss, K., Gentry, J. & Van der Auwera, G. Full-stack genomics pipelining with GATK4 + WDL + Cromwell. in (F1000 Research, 2017). doi:10.7490/f1000research.1114631.1.

49. Beauchamp, E. M. et al. ZBTB33 Is Mutated in Clonal Hematopoiesis and Myelodysplastic Syndromes and Impacts RNA Splicing. Blood Cancer Discov (2021) doi:10.1158/2643-3230.BCD-20-0224.

50. Miller, C. A. et al. Failure to detect mutations in U2AF1 due to changes in the GRCh38 reference sequence. 2021.05.07.442430 https://www.biorxiv.org/content/10.1101/2021.05.07.442430v1 (2021) doi:10.1101/2021.05.07.442430.

51. Hiatt, J. B., Pritchard, C. C., Salipante, S. J., O’Roak, B. J. & Shendure, J. Single molecule molecular inversion probes for targeted, high-accuracy detection of low-frequency variation. Genome Res. 23, 843–854 (2013).

52. mimips. (kitzmanlab, 2020).

53. Koboldt, D. C. et al. VarScan 2: Somatic mutation and copy number alteration discovery in cancer by exome sequencing. Genome Res. 22, 568–576 (2012).

54. Robinson, J. T. et al. Integrative genomics viewer. Nature Biotechnology 29, 24–26 (2011).

55. Ma, C., Blackwell, T., Boehnke, M. & Scott, L. J. Recommended Joint and Meta-Analysis Strategies for Case-Control Association Testing of Single Low-Count Variants. Genetic Epidemiology 37, 539–550 (2013).

56. Li, Z. et al. Dynamic Scan Procedure for Detecting Rare-Variant Association Regions in Whole-Genome Sequencing Studies. The American Journal of Human Genetics 104, 802–814 (2019).

57. Pedersen, B. S. & Quinlan, A. R. cyvcf2: fast, flexible variant analysis with Python. Bioinformatics 33, 1867–1869 (2017).

58. Bates, D. et al. Matrix: Sparse and Dense Matrix Classes and Methods. (2019).

59. R Core Team. R: A Language and environment for statistical computing. (R Foundation for Statistical Computing, 2020).

60. Gogarten, S. M. et al. Genetic association testing using the GENESIS R/Bioconductor package. Bioinformatics 35, 5346–5348 (2019).

61. W. N. Venables & B. D. Ripley. Modern Applied Statistics with S. (Springer, 2002).

62. Stan Development Team. Stan Modeling Language Users Guide and Reference Manual, 2.17. (2020).

63. Stan Development Team. RStan: The R interface to Stan. (2020).

64. Bezanson, J., Edelman, A., Karpinski, S. & Shah, V. B. Julia: A fresh approach to numerical computing. (2017).

65. Corces, M. R. et al. Lineage-specific and single-cell chromatin accessibility charts human hematopoiesis and leukemia evolution. Nature Genetics 48, 1193–1203 (2016).

66. Weber, E. W. et al. Transient “rest” restores functionality in exhausted CAR-T cells via epigenetic remodeling. Science 372, eaba1786 (2021).

67. Omni-ATAC-seq: Improved ATAC-seq protocol. https://www.researchsquare.com (2017) doi:10.1038/protex.2017.096.

68. Giambartolomei, C. et al. Bayesian Test for Colocalisation between Pairs of Genetic Association Studies Using Summary Statistics. PLOS Genetics 10, e1004383 (2014).

