## Supplementary Acknowledgements for "Clonal hematopoiesis is driven by aberrant activation of TCL1A"

**African-American Coronary Artery Calcium consortium study**

MESA Family is conducted and supported by the National Heart, Lung, and Blood Institute (NHLBI) in collaboration with MESA investigators. Support is provided by grants and contracts R01HL071051, R01HL071205, R01HL071250, R01HL071251, R01HL071258, R01HL071259, by the National Center for Research Resources, Grant UL1RR033176. Also supported in part by the National Center for Advancing Translational Sciences, CTSI grant UL1TR001881, and the National Institute of Diabetes and Digestive and Kidney Disease Diabetes Research Center (DRC) grant DK063491 to the Southern California Diabetes Endocrinology Research Center.

**Atrial Fibrillation Biobank Ludwig Maximilian University (AFLMU)**

Whole genome sequencing (WGS) for the Trans-Omics in Precision Medicine (TOPMed) program was supported by the National Heart, Lung and Blood Institute (NHLBI). WGS for “NHLBI TOPMed: Defining time-dependent genetic and transcriptomic responses to cardiac injury among patients with arrhythmias” (phs001543) was performed at the Broad Institute of MIT and Harvard (UM1HG008895). This funding source was an NHLBI supplement to NHGRI's Centers for Common Disease Genomics (CCDG). Centralized read mapping and genotype calling, along with variant quality metrics and filtering were provided by the TOPMed Informatics Research Center (3R01HL-117626-02S1). Phenotype harmonization, data management, sample-identity QC, and general study coordination, were provided by the TOPMed Data Coordinating Center (3R01HL-120393-02S1). We gratefully acknowledge the studies and participants who provided biological samples and data for TOPMed.

This study is part of the Centers for Common Disease Genomics (CCDG) program, a large-scale genome sequencing effort to identify rare risk and protective alleles that contribute to a range of common disease phenotypes. The CCDG program is funded by the National Human Genome Research Institute (NHGRI) and the National Heart, Lung, and Blood Institute (NHLBI). Sequencing was completed at the Human Genome Sequencing Center at Baylor College of Medicine under NHGRI grant UM1 HG008898. The cohort participants were recruited at the University Hospital of the LMU Munich, Munich, Germany, additionally facilitated through past funding by the German Federal Ministry of Research (BMBF) and the German Competence Network on Atrial Fibrillation (AFNET).

**Genetics of Cardiometabolic Health in the Amish**

The TOPMed component of the Amish Research Program was supported by NIH grants R01 HL121007, U01 HL072515, and R01 AG18728.

**Atherosclerosis Risk in Communities**

The Atherosclerosis Risk in Communities study has been funded in whole or in part with Federal funds from the National Heart, Lung, and Blood Institute, National Institutes of Health, Department of Health and Human Services (contract numbers HHSN268201700001I, HHSN268201700002I, HHSN268201700003I, HHSN268201700004I and HHSN268201700005I). The authors thank the staff and participants of the ARIC study for their important contributions.

**Australian Familial Atrial Fibrillation Study**

The NHGRI supplements were awarded with funds from the NHLBI.

**Barbados Asthma Genetics Study**

We gratefully acknowledge the contributions of Pissamai and Trevor Maul, Paul Levett, Anselm Hennis, P. Michele Lashley, Raana Naidu, Malcolm Howitt and Timothy Roach, and the numerous health care providers, and community clinics and co-investigators who assisted in the phenotyping and collection of DNA samples, and the families and patients for generously donating DNA samples to the Barbados Asthma Genetics Study (BAGS). Funding for BAGS was provided by National Institutes of Health (NIH) R01HL104608, R01HL087699, and HL104608 S1.

**Mount Sinai BioMe Biobank**

The Mount Sinai BioMe Biobank has been supported by The Andrea and Charles Bronfman Philanthropies and in part by Federal funds from the NHLBI and NHGRI (U01HG00638001; U01HG007417; X01HL134588). We thank all participants in the Mount Sinai Biobank. We also thank all our recruiters who have assisted and continue to assist in data collection and management and are grateful for the computational resources and staff expertise provided by Scientific Computing at the Icahn School of Medicine at Mount Sinai.

**Duke Cardiac CATHeterization (CATHGEN) study**

Funding support for the Genetic Mediators of Metabolic CVD Risk was provided by NHLBI grant RC2 HL101621 (William E. Kraus)

**The Cleveland Clinic Atrial Fibrillation Study**

The research reported in this article was supported by grants from the National Institutes of Health (NIH) National Heart, Lung, and Blood Institute grants R01 HL090620 and R01 HL111314, the NIH National Center for Research Resources for Case Western Reserve University and Cleveland Clinic Clinical and Translational Science Award (CTSA) UL1-RR024989, the Department of Cardiovascular Medicine philanthropic research fund, Heart and Vascular Institute, Cleveland Clinic, the Fondation Leducq grant 07-CVD 03, and The Atrial Fibrillation Innovation Center, State of Ohio.

**Whole Genome Sequence Analysis in Early Cerebral Small Vessel Disease**

The Coronary Artery Risk Development in Young Adults Study (CARDIA) is conducted and supported by the National Heart, Lung, and Blood Institute (NHLBI) in collaboration with the University of Alabama at Birmingham (HHSN268201800005I & HHSN268201800007I), Northwestern University (HHSN268201800003I), University of Minnesota (HHSN268201800006I), and Kaiser Foundation Research Institute (HHSN268201800004I). CARDIA was also partially supported by the Intramural Research Program of the National Institute on Aging (NIA) and an intra‐agency agreement between NIA and NHLBI (AG0005).

**Cleveland Family Study**

The Cleveland Family Study has been supported in part by National Institutes of Health grants [R01-HL046380, KL2-RR024990, R35-HL135818, and R01-HL113338].

**Cardiovascular Health Study**

Cardiovascular Health Study: This research was supported by contracts HHSN268201200036C, HHSN268200800007C, HHSN268201800001C, N01HC55222, N01HC85079, N01HC85080, N01HC85081, N01HC85082, N01HC85083, N01HC85086, 75N92021D00006, and grants U01HL080295 and U01HL130114 from the National Heart, Lung, and Blood Institute (NHLBI), with additional contribution from the National Institute of Neurological Disorders and Stroke (NINDS). Additional support was provided by R01AG023629 from the National Institute on Aging (NIA). A full list of principal CHS investigators and institutions can be found at CHS-NHLBI.org. The content is solely the responsibility of the authors and does not necessarily represent the official views of the National Institutes of Health.

**Genetic Epidemiology of COPD**

The COPDGene project described was supported by Award Number U01 HL089897 and Award Number U01 HL089856 from the National Heart, Lung, and Blood Institute. The content is solely the responsibility of the authors and does not necessarily represent the official views of the National Heart, Lung, and Blood Institute or the National Institutes of Health. The COPDGene project is also supported by the COPD Foundation through contributions made to an Industry Advisory Board that has included AstraZeneca, Bayer Pharmaceuticals, Boehringer Ingelheim, Genentech, GlaxoSmithKline, Novartis, Pfizer, and Sunovion. A full listing of COPDGene investigators can be found at: http://www.copdgene.org/directory

**Determining the association of chromosomal variants with non-PV triggers and ablation-outcome in AF**

Whole genome sequencing (WGS) for the Trans-Omics in Precision Medicine (TOPMed) program was supported by the National Heart, Lung and Blood Institute (NHLBI). The TOPMed acknowledgement statement can be found at: https://www.nhlbiwgs.org/acknowledgements. WGS for “NHLBI TOPMed: Determining the association of chromosomal variants with non-PV triggers and ablation-outcome in AF-DECAF” (phs001546.v1.p1) was performed at the Broad Institute of MIT and Harvard (3U54HG003067-12S2, 3U54HG003067-13S1). Centralized read mapping and genotype calling, along with variant quality metrics and filtering were provided by the TOPMed Informatics Research Center (3R01HL-117626-02S1). Phenotype harmonization, data management, sample-identity QC, and general study coordination, were provided by the TOPMed Data Coordinating Center (3R01HL-120393-02S1). We gratefully acknowledge the studies and participants who provided biological samples and data for TOPMed. The NHGRI supplements were awarded with funds from the NHLBI.

**Diabetes Heart Study**

This work was supported by R01 HL92301, R01 HL67348, R01 NS058700, R01 AR48797, R01 DK071891, R01 AG058921, the General Clinical Research Center of the Wake Forest University School of Medicine (M01 RR07122, F32 HL085989), the American Diabetes Association, and a pilot grant from the Claude Pepper Older Americans Independence Center of Wake Forest University Health Sciences (P60 AG10484).

**Early-onset Atrial Fibrillation in the Estonian Biobank**

The TOPMed acknowledgement statement can be found at: https://www.nhlbiwgs.org/acknowledgements. Please include: "Whole genome sequencing (WGS) for the Trans-Omics in Precision Medicine (TOPMed) program was supported by the National Heart, Lung and Blood Institute (NHLBI). WGS for “NHLBI TOPMed: Defining time-dependent genetic and transcriptomic responses to cardiac injury among patients with arrhythmias” (phs# with version extension) was performed at the Broad Institute of MIT and Harvard (3R01HL092577-06S1). Centralized read mapping and genotype calling, along with variant quality metrics and filtering were provided by the TOPMed Informatics Research Center (3R01HL-117626-02S1). Phenotype harmonization, data management, sample-identity QC, and general study coordination, were provided by the TOPMed Data Coordinating Center (3R01HL-120393-02S1). We gratefully acknowledge the studies and participants who provided biological samples and data for TOPMed."

**Evaluation of COPD Longitudinally to Identify Predictive Surrogate Endpoints**

The ECLIPSE study (NCT00292552) was sponsored by GlaxoSmithKline. The ECLIPSE investigators included: ECLIPSE Investigators — Bulgaria: Y. Ivanov, Pleven; K. Kostov, Sofia. Canada: J. Bourbeau, Montreal; M. Fitzgerald, Vancouver, BC; P. Hernandez, Halifax, NS; K. Killian, Hamilton, ON; R. Levy, Vancouver, BC; F. Maltais, Montreal; D. O'Donnell, Kingston, ON. Czech Republic: J. Krepelka, Prague. Denmark: J. Vestbo, Hvidovre. The Netherlands: E. Wouters, Horn-Maastricht. New Zealand: D. Quinn, Wellington. Norway: P. Bakke, Bergen. Slovenia: M. Kosnik, Golnik. Spain: A. Agusti, J. Sauleda, P. de Mallorca. Ukraine: Y. Feschenko, V. Gavrisyuk, L. Yashina, Kiev; N. Monogarova, Donetsk. United Kingdom: P. Calverley, Liverpool; D. Lomas, Cambridge; W. MacNee, Edinburgh; D. Singh, Manchester; J. Wedzicha, London. United States: A. Anzueto, San Antonio, TX; S. Braman, Providence, RI; R. Casaburi, Torrance CA; B. Celli, Boston; G. Giessel, Richmond, VA; M. Gotfried, Phoenix, AZ; G. Greenwald, Rancho Mirage, CA; N. Hanania, Houston; D. Mahler, Lebanon, NH; B. Make, Denver; S. Rennard, Omaha, NE; C. Rochester, New Haven, CT; P. Scanlon, Rochester, MN; D. Schuller, Omaha, NE; F. Sciurba, Pittsburgh; A. Sharafkhaneh, Houston; T. Siler, St. Charles, MO; E. Silverman, Boston; A. Wanner, Miami; R. Wise, Baltimore; R. ZuWallack, Hartford, CT. ECLIPSE Steering Committee: H. Coxson (Canada), C. Crim (GlaxoSmithKline, USA), L. Edwards (GlaxoSmithKline, USA), D. Lomas (UK), W. MacNee (UK), E. Silverman (USA), R. Tal-Singer (Co-chair, GlaxoSmithKline, USA), J. Vestbo (Co-chair, Denmark), J. Yates (GlaxoSmithKline, USA). ECLIPSE Scientific Committee: A. Agusti (Spain), P. Calverley (UK), B. Celli (USA), C. Crim (GlaxoSmithKline, USA), B. Miller (GlaxoSmithKline, USA), W. MacNee (Chair, UK), S. Rennard (USA), R. Tal-Singer (GlaxoSmithKline, USA), E. Wouters (The Netherlands), J. Yates (GlaxoSmithKline, USA).

**Boston Early-Onset COPD Study**

The Boston Early-Onset COPD Study was supported by R01 HL113264 and U01 HL089856 from the National Heart, Lung, and Blood Institute.

**Framingham Heart Study**

The Framingham Heart Study (FHS) acknowledges the support of contracts NO1-HC-25195, HHSN268201500001I and 75N92019D00031 from the National Heart, Lung and Blood Institute and grant supplement R01 HL092577-06S1 for this research. We also acknowledge the dedication of the FHS study participants without whom this research would not be possible. Dr. Vasan is supported in part by the Evans Medical Foundation and the Jay and Louis Coffman Endowment from the Department of Medicine, Boston University School of Medicine.

**Genetic Epidemiology Network of Arteriopathy**

Support for GENOA was provided by the National Heart, Lung and Blood Institute (HL054457, HL054464, HL054481, HL119443, and HL087660) of the National Institutes of Health. GENOA may also require study-specific acknowledgements depending on the phenotypes analyzed (please consult with the GENOA co-author on the paper).

**Genetic Epidemiology Network of Salt Sensitivity**

The Genetic Epidemiology Network of Salt-Sensitivity (GenSalt) was supported by research grants (U01HL072507, R01HL087263, and R01HL090682) from the National Heart, Lung and Blood Institute, National Institutes of Health, Bethesda, MD.

**GeneSTAR**

GeneSTAR was supported by the National Institutes of Health/National Heart, Lung, and Blood Institute (U01 HL72518, HL087698, HL112064) and by a grant from the National Institutes of Health/National Center for Research Resources (M01-RR000052) to the Johns Hopkins General Clinical Research Center.

**Groningen Genetics of Atrial Fibrillation**

The TOPMed acknowledgement statement can be found at: https://www.nhlbiwgs.org/acknowledgements. In addition, please include: "Whole genome sequencing (WGS) for the Trans-Omics in Precision Medicine (TOPMed) program was supported by the National Heart, Lung and Blood Institute (NHLBI). WGS for “NHLBI TOPMed - Centers for Common Disease Genomics (CCDG): Groningen Genetics of Atrial Fibrillation (GGAF)” (phs# with version extension) was performed at the Baylor College of Medicine. This funding source was an NHLBI supplement to NHGRI's Centers for Common Disease Genomics (CCDG). Centralized read mapping and genotype calling, along with variant quality metrics and filtering were provided by the TOPMed Informatics Research Center (3R01HL-117626-02S1). Phenotype harmonization, data management, sample-identity QC, and general study coordination, were provided by the TOPMed Data Coordinating Center (3R01HL-120393-02S1). We gratefully acknowledge the studies and participants who provided biological samples and data for TOPMed."

**Genetics of Lipid Lowering Drugs and Diet Network**

GOLDN biospecimens, baseline phenotype data, and intervention phenotype data were collected with funding from National Heart, Lung and Blood Institute (NHLBI) grant U01 HL072524. Whole-genome sequencing in GOLDN was funded by NHLBI grant R01 HL104135 and supplement R01 HL104135-04S1.

**Hispanic Community Health Study - Study of Latinos**

The Hispanic Community Health Study/Study of Latinos is a collaborative study supported by contracts from the National Heart, Lung, and Blood Institute (NHLBI) to the University of North Carolina (HHSN268201300001I / N01-HC-65233), University of Miami (HHSN268201300004I / N01-HC-65234), Albert Einstein College of Medicine (HHSN268201300002I / N01-HC-65235), University of Illinois at Chicago – HHSN268201300003I / N01-HC-65236 Northwestern Univ), and San Diego State University (HHSN268201300005I / N01-HC-65237). The following Institutes/Centers/Offices have contributed to the HCHS/SOL through a transfer of funds to the NHLBI: National Institute on Minority Health and Health Disparities, National Institute on Deafness and Other Communication Disorders, National Institute of Dental and Craniofacial Research, National Institute of Diabetes and Digestive and Kidney Diseases, National Institute of Neurological Disorders and Stroke, NIH Institution-Office of Dietary Supplements.

**Heart and Vascular Health Study**

The Heart and Vascular Health Study was supported by grants HL068986, HL085251, HL095080, and HL073410 from the National Heart, Lung, and Blood Institute.

**Hypertension Genetic Epidemiology Network**

The HyperGEN Study is part of the National Heart, Lung, and Blood Institute (NHLBI) Family Blood Pressure Program; collection of the data represented here was supported by grants U01 HL054472 (MN Lab), U01 HL054473 (DCC), U01 HL054495 (AL FC), and U01 HL054509 (NC FC). The HyperGEN: Genetics of Left Ventricular Hypertrophy Study was supported by NHLBI grant R01 HL055673 with whole-genome sequencing made possible by supplement -18S1.

**Jackson Heart Study**

The Jackson Heart Study (JHS) is supported and conducted in collaboration with Jackson State University (HHSN268201800013I), Tougaloo College (HHSN268201800014I), the Mississippi State Department of Health (HHSN268201800015I) and the University of Mississippi Medical Center (HHSN268201800010I, HHSN268201800011I and HHSN268201800012I) contracts from the National Heart, Lung, and Blood Institute (NHLBI) and the National Institute on Minority Health and Health Disparities (NIMHD). The authors also wish to thank the staffs and participants of the JHS.

The views expressed in this manuscript are those of the authors and do not necessarily represent the views of the National Heart, Lung, and Blood Institute; the National Institutes of Health; or the U.S. Department of Health and Human Services.

**Mayo Clinic Venous Thromboembolism Study**

Funded, in part, by grants from the National Institutes of Health, National Heart, Lung and Blood Institute (HL66216 and HL83141). the National Human Genome Research Institute (HG04735, HG06379), and research support provided by Mayo Foundation.

**Multi-Ethnic Study of Atherosclerosis**

The MESA project is conducted and supported by the National Heart, Lung, and Blood Institute (NHLBI) in collaboration with MESA investigators. Support for MESA is provided by contracts 75N92020D00001, HHSN268201500003I, N01-HC-95159, 75N92020D00005, N01-HC-95160, 75N92020D00002, N01-HC-95161, 75N92020D00003, N01-HC-95162, 75N92020D00006, N01-HC-95163, 75N92020D00004, N01-HC-95164, 75N92020D00007, N01-HC-95165, N01-HC-95166, N01-HC-95167, N01-HC-95168, N01-HC-95169, UL1-TR-000040, UL1-TR-001079, UL1-TR-001420. Also supported in part by the National Center for Advancing Translational Sciences, CTSI grant UL1TR001881, and the National Institute of Diabetes and Digestive and Kidney Disease Diabetes Research Center (DRC) grant DK063491 to the Southern California Diabetes Endocrinology Research Center. We gratefully acknowledge the studies and participants who provided biologic samples and data for MESA and TOPMed.

**Massachusetts General Hospital (MGH) Atrial Fibrillation Study**

The research reported in this article was supported by NIH grants K23HL071632, K23HL114724, R21DA027021, R01HL092577, R01HL092577S1, R01HL104156, K24HL105780, and U01HL65962. The research has also been supported by an Established Investigator Award from the American Heart Association (13EIA14220013) and by support from the Fondation Leducq (14CVD01). This manuscript was not prepared in collaboration with MGH AF Study investigators and does not necessarily reflect the opinions or views of the MGH AF Study investigators or the NHLBI.

**Defining the time-dependent genetic and transcriptomic responses to cardiac injury among patients with arrhythmias**

The NHGRI supplements were awarded with funds from the NHLBI.

**My Life, Our Future: Genotyping for Progress in Hemophilia**

The My Life, Our Future samples and data are made possible through the partnership of Bloodworks Northwest, the American Thrombosis and Hemostasis Network, the National Hemophilia Foundation, and Bioverativ. We gratefully acknowledge the hemophilia treatment centers and their patients who provided biological samples and phenotypic data.

**Malmo Preventive Project**

Whole genome sequencing (WGS) for the Trans-Omics in Precision Medicine (TOPMed) program was supported by the National Heart, Lung and Blood Institute (NHLBI). WGS for “NHLBI TOPMed: Defining time-dependent genetic and transcriptomic responses to cardiac injury among patients with arrhythmias” (phs001544) was performed at the Broad Institute of MIT and Harvard (UM1HG008895). This funding source was an NHLBI supplement to NHGRI's Centers for Common Disease Genomics (CCDG). Centralized read mapping and genotype calling, along with variant quality metrics and filtering were provided by the TOPMed Informatics Research Center (3R01HL-117626-02S1). Phenotype harmonization, data management, sample-identity QC, and general study coordination, were provided by the TOPMed Data Coordinating Center (3R01HL-120393-02S1). We gratefully acknowledge the studies and participants who provided biological samples and data for TOPMed. This study is part of the Centers for Common Disease Genomics (CCDG) program, a large-scale genome sequencing effort to identify rare risk and protective alleles that contribute to a range of common disease phenotypes. The CCDG program is funded by the National Human Genome Research Institute (NHGRI) and the National Heart, Lung, and Blood Institute (NHLBI). Sequencing was completed at the Human Genome Sequencing Center at Baylor College of Medicine under NHGRI grant UM1 HG008898.

**Partners HealthCare Biobank**

We thank the Broad Institute for generating high-quality sequence data supported by the NHLBI grant 3R01HL092577-06S1 to Dr. Patrick Ellinor

**San Antonio Family Studies**

Collection of the San Antonio Family Study data was supported in part by National Institutes of Health (NIH) grants R01 HL045522, MH078143, MH078111 and MH083824; and whole genome sequencing of SAFS subjects was supported by U01 DK085524 and R01 HL113323. We are very grateful to the participants of the San Antonio Family Study for their continued involvement in our research programs.

**Study of Asthma Phenotypes and Pharmacogenomic Interactions by Race-Ethnicity**

Grant funding sources must be acknowledged in any study using SAPPHIRE data. These include the Fund for Henry Ford Hospital, the American Asthma Foundation, the National Heart Lung and Blood Institute (R01HL118267, X01HL134589), the National Institute of Allergy and Infectious Diseases (R01AI079139), and the National Institute of Diabetes and Digestive and Kidney Diseases (R01DK113003).

**Study of Adiposity in Samoans**

Date collection was funded by NIH grant R01-HL093093. We thank the Samoan participants of the study and local village authorities. We acknowledge the support of the Samoan Ministry of Health and the Samoa Bureau of Statistics for their support of this research.

**Taiwan Study of Hypertension using Rare Variants**

The Rare Variants for Hypertension in Taiwan Chinese (THRV) is supported by the National Heart, Lung, and Blood Institute (NHLBI) grant (R01HL111249) and its participation in TOPMed is supported by an NHLBI supplement (R01HL111249-04S1). THRV is a collaborative study between Washington University in St. Louis, LA BioMed at Harbor UCLA, University of Texas in Houston, Taichung Veterans General Hospital, Taipei Veterans General Hospital, Tri-Service General Hospital, National Health Research Institutes, National Taiwan University, and Baylor University. THRV is based (substantially) on the parent SAPPHIRe study, along with additional population-based and hospital-based cohorts. SAPPHIRe was supported by NHLBI grants (U01HL54527, U01HL54498) and Taiwan funds, and the other cohorts were supported by Taiwan funds.

**The Women's Genome Health Study**

The WGHS is supported by the National Heart, Lung, and Blood Institute (HL043851 and HL080467) and the National Cancer Institute (CA047988 and UM1CA182913). The most recent cardiovascular endpoints were supported by ARRA funding HL099355.

**Women's Health Initiative**

The WHI program is funded by the National Heart, Lung, and Blood Institute, National Institutes of Health, U.S. Department of Health and Human Services through contracts 75N92021D00001, 75N92021D00002, 75N92021D00003, 75N92021D00004, 75N92021D00005.
