## Supplementary Theoretical Text for "Clonal hematopoiesis is driven by aberrant activation of TCL1A"

HSC simulation results

### PACER Simulation Parameters

We assume that the accumulation of passenger mutations is described by a Poisson birth-death stochastic process. As the birth and death rates scale with the number of HSCs, we assume a linear birth-death process.

We assume that the birth rate for a given hematopoietic stem cell (HSC) $i$ at time $t$ with fitness $s_{i}(t)$) is $\lambda_{i}(t)\sim Poisson(\omega*X_{i}(t)*(1+s_{i}(t))*dt)$, where $dt$ represents the amount of time in years, and $\omega$ represents the number of stem cell divisions per year. We assume that the death rate can be described as

$$\psi_{i}(t)\sim Poisson(\omega*X_{i}(t)*(1-s_{i}(t))*dt)$$

. The death rate is the rate at which an HSC divides into two differentiated cells, and the birth rate is the rate at which an HSC divides into two HSCs. We don’t consider asymmetric HSC differentiation as this would not change the clone size. The HSC clone cell count is defined as $X_{i}(t)=\sum_{l\leq t} \lambda_{i}(l)-\psi_{i}(l)$, and the HSC clone size (a fraction of the total cell population) is $VAF_{i}(t)=\frac{X_{i}(t)}{\sum_{j} X_{j}(t)}$.

We start with 500 HSC clones, each with 200 identical cells in each clone $X_{i}(t=0)=200$. Each cell divides once every three years (= 1/3), and each clone with an initial $s_{i}(t=0)=0$. At each iteration, we also center the $s_{i}(t)$ such that $\overline{s_{i}(t)}=0$. This means that there are 100,000 total HSCs at the start of the simulation.

For each clone, we set the passenger mutation rate:

1. $\mu_{p}$, the passenger accumulation rate, $A_{i}(t)\sim Poisson(X_{i}(t)*\mu_{p}*dt)$

Where $A_{i}(t)$ is the number of passengers accumulated in a given clone through time $t$. We set $\mu_{p}=0.006$, which is the passenger mutation rate of a diploid genome for a single HSC per year. This implies a mutation rate of 6 passengers per year for a clone with 1000 cells, and a mutation rate of 600 passengers per year across the entire population of 100,000 HSCs. We will later consider the effects of an insensitive sequencing assay that captures a small fraction of the passengers.

We assign a single driver to one of the HSC clones, which is randomly selected among the HSC clones. The time of acquisition is uniformly drawn from each cell division after 10 years, such that are driver is equally likely to be acquired at either 10 years or 78 years. We simulate the HSC population across a lifetime of 90 years. We refer to the time of driver acquision as $T_{d}$.

We assume that each HSC clone can at most acquire a single driver, which represents a similar HSC population to the TOPMed CHIP driver carriers.

If an HSC clone $i$ acquires a driver at time $t$, we set $s_{i}(t)=Beta(4,16)$. A $Beta(4,16)$ random variable is bounded between 0 and 1 and has an expectation of 0.20. An HSC with ${}_{i}(t)=0.20$ will self-renew 60% of the time, and terminally differentiate 40% of the time.

For a given HSC population, we simulate 90 years, and track the accumulation of passengers and drivers. To incorporate the censoring from using 38x sequencing coverage, we simulate whether a given passenger would be observed at 38x coverage by sampling the number of alt-reads from $R\sim Binomial(38,VAF_{i}(t))$ and comparing $R\geq2$, since two reads are required by our variant calling process. We refer to $P(R\geq2|VAF=vaf)=P(Binomial(38,vaf)\geq2)$ We refer to the passengers that would be detected at 38x coverage as the censored founding passengers, $AC_{i}(t)$, where $t=T_{d}$ .

We ran the simulation 10,000 times, where at most a single HSC clone acquires a driver mutation. We then compared $AC_{i}(T_{d})$ to the fitness of the clone at the end of each simulation.

### PACER Simulation Results

#### Supplementary Figure S6

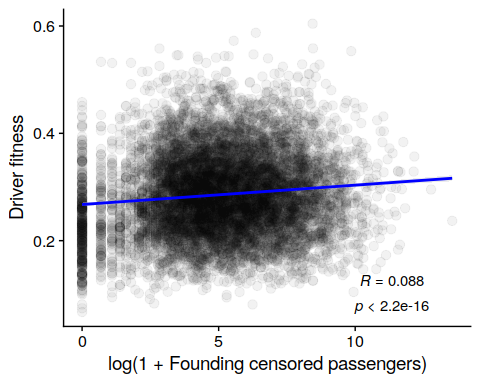

We observed a modest concordance between the founding censored passengers with the fitness of each HSC at the end of the simulation (spearman = 0.09, pvalue < 2.2e-16, Supplementary Figure S6). This suggests that the censored founding passengers are proportional to the fitness of the driver mutations. Stochastic drift of the HSC clone sizes contributes substantial residual variance.

#### Supplementary Figure S7

We observe no concordance between the uncensored founding passengers with fitness (Supplementary Figure S7).

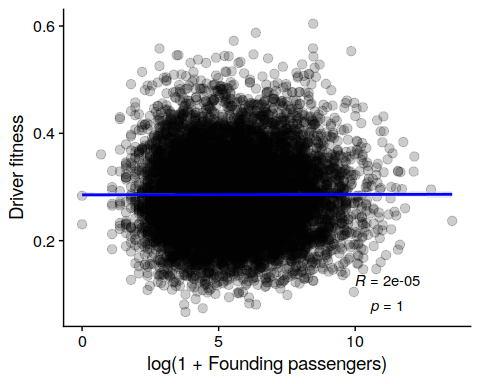
