## Supplementary figures and images for "Clonal hematopoiesis is driven by aberrant activation of TCL1A"

### Supplementary Figure 1

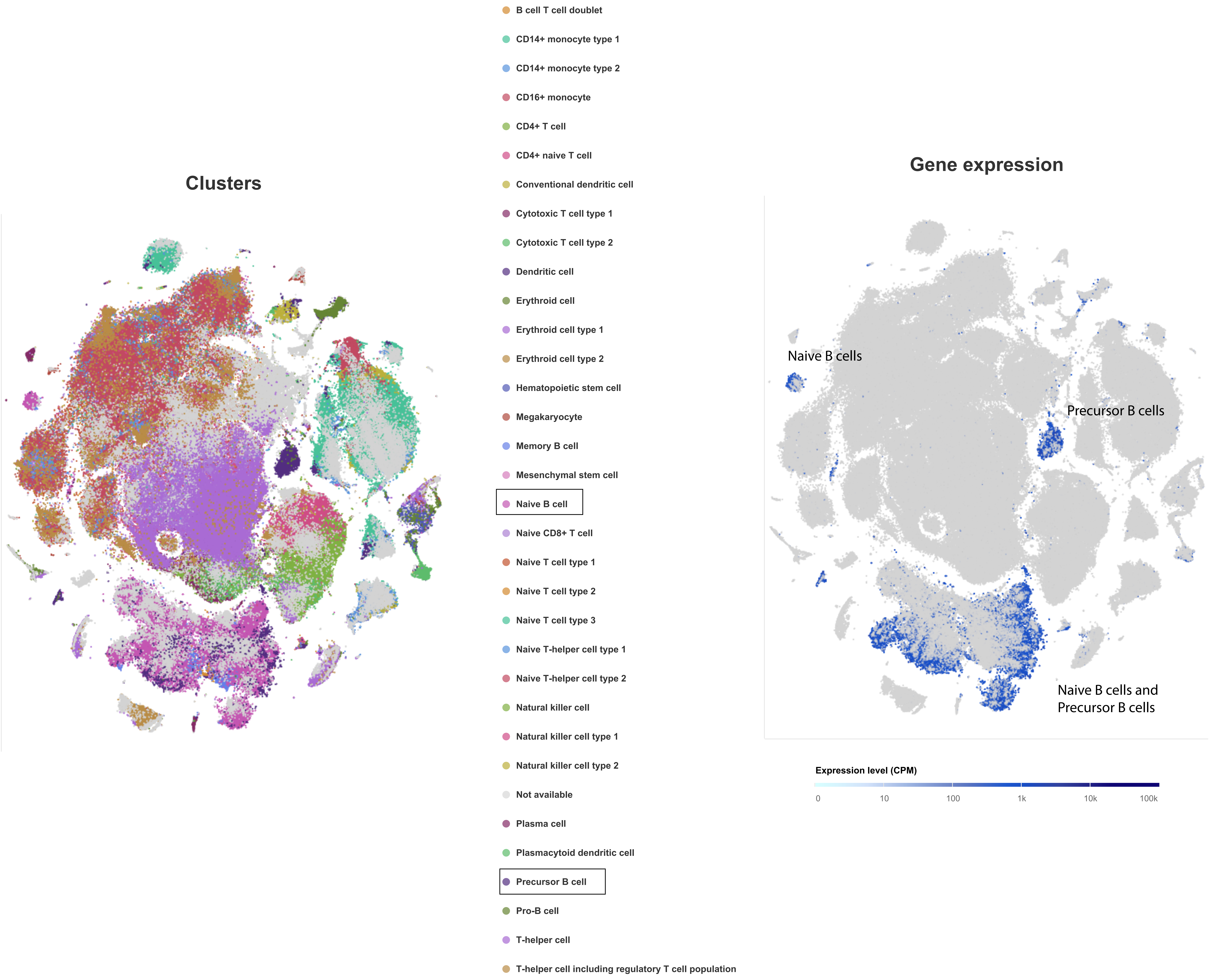
